# Vgll3a promotes sexual maturation in male and female Atlantic salmon

**DOI:** 10.64898/2026.06.19.733363

**Authors:** Erik Kjærner-Semb, Thomas W. K. Fraser, Petra Vogelsang, Kai Skaftnesmo, Fernando Ayllon, Rolf B. Edvardsen, Sebastian Braathen, Birgitta Norberg, Per Gunnar Fjelldal, Eva Andersson, Rüdiger W. Schulz, Anna Wargelius

## Abstract

The age at which Atlantic salmon reaches sexual maturity shows a strong hereditary component associated with the *vgll3a* locus. The role of Vgll3 in maturation has remained unknown in vertebrates until recently, when it has been linked to pleiotropic roles in killifish, both delaying male maturation and affecting lifespan by protecting against cancer. As Atlantic salmon has two *vgll3* paralogs, where only *vgll3a* has been associated with sexual maturation, it may provide a suitable model for studying the maturation-specific function of *vgll3*, as the other paralog may buffer for pleiotropic roles of *vgll3*. To address this, we used CRISPR/Cas9 to generate fish highly mutated in the *vgll3a* gene. We monitored their maturation and crossed highly mutated crispants to generate two year-classes of complete loss-of-function. All groups were reared under environmental conditions triggering early maturation in one-year-old males. We found a clear difference in the proportion of sexually maturing or mature fish between the different genotypes: in all experiments significantly fewer *vgll3a^-/-^* males entered puberty and reached final maturation compared to *vgll3a^+/-^* and *vgll3a^+/+^* males. Furthermore, loss of *vgll3a* resulted in lower frequencies of maturation also in females. We conclude that Vgll3a stimulates maturation and that its complete removal significantly reduced maturation rates in both sexes in Atlantic salmon. Our findings also identify *vgll3a* as the causative gene in the locus associated with age at sexual maturity. Together, our findings support a new role for Vgll3 in initiating puberty in vertebrates and identifying salmon as a promising model for functional studies regarding the timing of sexual maturation.

## 1. Introduction

A major developmental role of Vgll3 has been established in teleost fish, where it has been linked to the timing of sexual maturation. In Atlantic salmon (*Salmo salar*), genome-wide association studies identified *vgll3a* as the locus with the strongest effect on age at maturity, explaining over 30% of the phenotypic variance in both males and females (1,2). Allelic variation in *vgll3a* underpins differences in the propensity to mature early or late, supporting a model in which *vgll3a* influences timing of maturation. However, no study so far has linked the actual gene product to its function in maturation.

The Vestigial-like (VGLL) family of transcriptional cofactors traces its origin to the *Drosophila melanogaster* gene *vestigial* (*vg*), in which the encoded protein together with the DNA-binding protein Scalloped (Sd), is essential for wing development. Mammalian vestigial-like orthologs act as cofactors capable of interacting with the mammalian Sd orthologs TEAD/TEF-1 (3). Across bilaterians, the Vg/VGLL TEAD module is conserved: *Drosophila* encodes a single *vg* gene and a single TEAD ortholog (Sd), whereas vertebrates expanded the cofactor set to VGLL1–4 and the TEAD partners to TEAD1–4 (4,5). In invertebrates, the Vg Tondu (TDU) motif can convert a TEAD factor into a tissue-specific transcriptional activator (4,6,7). This functional logic is maintained in vertebrates, where VGLL proteins recruit TEADs to regulate lineage-specific gene expression programs in tissues such as muscle and placenta, thereby integrating Hippo-pathway signaling outputs (8). VGLL3 interacts primarily with TEAD1, TEAD3 and TEAD4 in muscle and other tissues, acting as a TEAD-dependent coactivator whose activity is modulated by developmental signals, mechanical cues and Hippo-pathway dynamics (9).

In wild European Atlantic salmon, age at maturation is strongly associated with the *vgll3a* locus on chromosome 25, with distinct alleles promoting either early (E) or late (L) maturation (1,2). Both alleles are present in most commercial salmon strains, and the L allele has shown promise in delaying male puberty in aquaculture (10,11). Owing to a salmonid-specific whole-genome duplication (12), Atlantic salmon encode two paralogs: *vgll3a* on chromosome 25 (associated with maturation timing) and *vgll3b* on chromosome 21. Both are expressed in male and female gonads, but only *vgll3a* is downregulated in the testis during puberty onset (13). Previous studies have mapped *vgll3a* expression to somatic support cells (Sertoli/granulosa) and linked it to Hippo signaling (13,14). Interestingly, allele-dependent changes in the expression of the pituitary hormone gene *fshb* were reported, with the E allele promoting higher expression than the L allele (15,16) . This aligns well with the observation that loss of the receptor for Fsh prevented puberty in salmon males (17) and supports the notion that Fsh is a crucial puberty inducing hormone in salmon.

In vertebrates, VGLL3 has roles across several biological processes. In mouse skeletal muscle, VGLL3 cooperates with TEAD1/3/4 to influence myogenic gene expression (18). Mechanosensitive functions of VGLL3 have also been described; in cardiac myofibroblasts, VGLL3 responds to matrix stiffness and drives profibrotic transcriptional programs and collagen production in mouse (19). In immunology, VGLL3 has emerged as a key regulator of female-biased autoimmune pathways: overexpression of VGLL3 in skin is sufficient to trigger cutaneous and systemic autoimmune disease resembling lupus, and it drives a gene network that contributes to the higher incidence of autoimmune disease in females (20). In cancer biology, VGLL3 can be amplified or overexpressed in subsets of sarcomas and other tumors, functioning as part of the TEAD-cofactor system, alongside YAP/TAZ and other VGLL paralogs, that influence proliferation and cell-state transitions (8). While genetic variation in the human VGLL3 locus has been associated with pubertal timing in females (21) the mechanistic role of VGLL3 in this process remains unclear. Recent work has demonstrated that Vgll3 is also associated with sexual maturation in killifish, while exhibiting pleiotropic effects on longevity. Specifically, males carrying a nonsense mutation in exon 1 displayed precocious maturation and developed melanoma at an earlier age compared with wild-type controls. In contrast, a nonsense mutation in exon 3 resulted in delayed male maturation, highlighting distinct functional effects of mutations in different regions of the gene (22). Salmon may provide a useful comparative model to investigate this function, considering that VGLL3 has been implicated in multiple biological processes in humans (8,20). After all, following genome duplication in salmonids, *vgll3* paralogs may have undergone sub-functionalization, potentially reducing the challenges associated with functional studies of pleiotropic genes.

Early male maturation poses a significant welfare and production challenge in salmon aquaculture, both in sea-cage and land-based recirculating aquaculture systems (RAS) (23,24). Precocious puberty increases vulnerability to disease and impairs osmoregulation, often leading to higher mortality, reduced growth and downgraded harvest quality. While manipulating photoperiod via artificial light can partly mitigate these problems, rising water temperatures, driven by climate change and increased use of closed systems, increase early maturation even under controlled light regimes (25). In general, Atlantic salmon females mature one or two years later than males, and precocious puberty in females is not a production problem. Instead, there is growing interest in identifying genetic factors that could shorten the generation time in females from three or four years to two, thereby accelerating genetic gain in breeding programs.

The overarching aim of this study was to determine how the *vgll3a* paralog regulates age at maturity in Atlantic salmon. We first generated highly mutated *vgll3a* crispants, monitored their maturation, and subsequently bred them to produce F1 progeny. From these offspring, we selected two year-classes representing the genotypes *vgll3a^-/-^*, *vgll3a^+/-^* and *vgll3a^+/+^*. All experimental groups were reared under environmental regimes known to induce precocious maturation in 1.5- to 2-year-old males, while female maturation was assessed at three and four years of age. Across all experiments, both sexes and multiple year-classes, the results were consistent in that *vgll3a^-/-^* and *vgll3a* crispant fish showed lower prevalence of maturation relative to *vgll3a^+/-^*, and *vgll3a^+/+^* individuals. Taken together, these findings support the conclusion that *vgll3a* promotes sexual maturation in Atlantic salmon males and females.

## 2. Results

### 2.1. Characterization of mutations in *vgll3a* crispants and F1 generations

The gene structure, CRISPR target site, and conservation of the *vgll3a* gene, as well as its CRISPR editing efficiency, are illustrated in Figure 1. The Vgll3a protein contains the conserved TONDU domain, which is located downstream of the CRISPR target site (Figure 1A). Of the two missense mutations previously linked to variation in age at maturation (1,2), one (T54M) is located upstream of this region, while the other (N323K) is located downstream. Amino acid sequence alignment showed strong conservation between Atlantic salmon and Northern pike, and moderate similarity to zebrafish and human VGLL3; however, low conservation was observed at the CRISPR target site (Figure 1A). Analysis of the *vgll3a* crispants revealed high total indel frequencies (Figure 1B) and a predominance of frameshift mutations over in-frame mutations in both males and females, indicating highly efficient and comparable editing across sexes.

**Figure 1.**
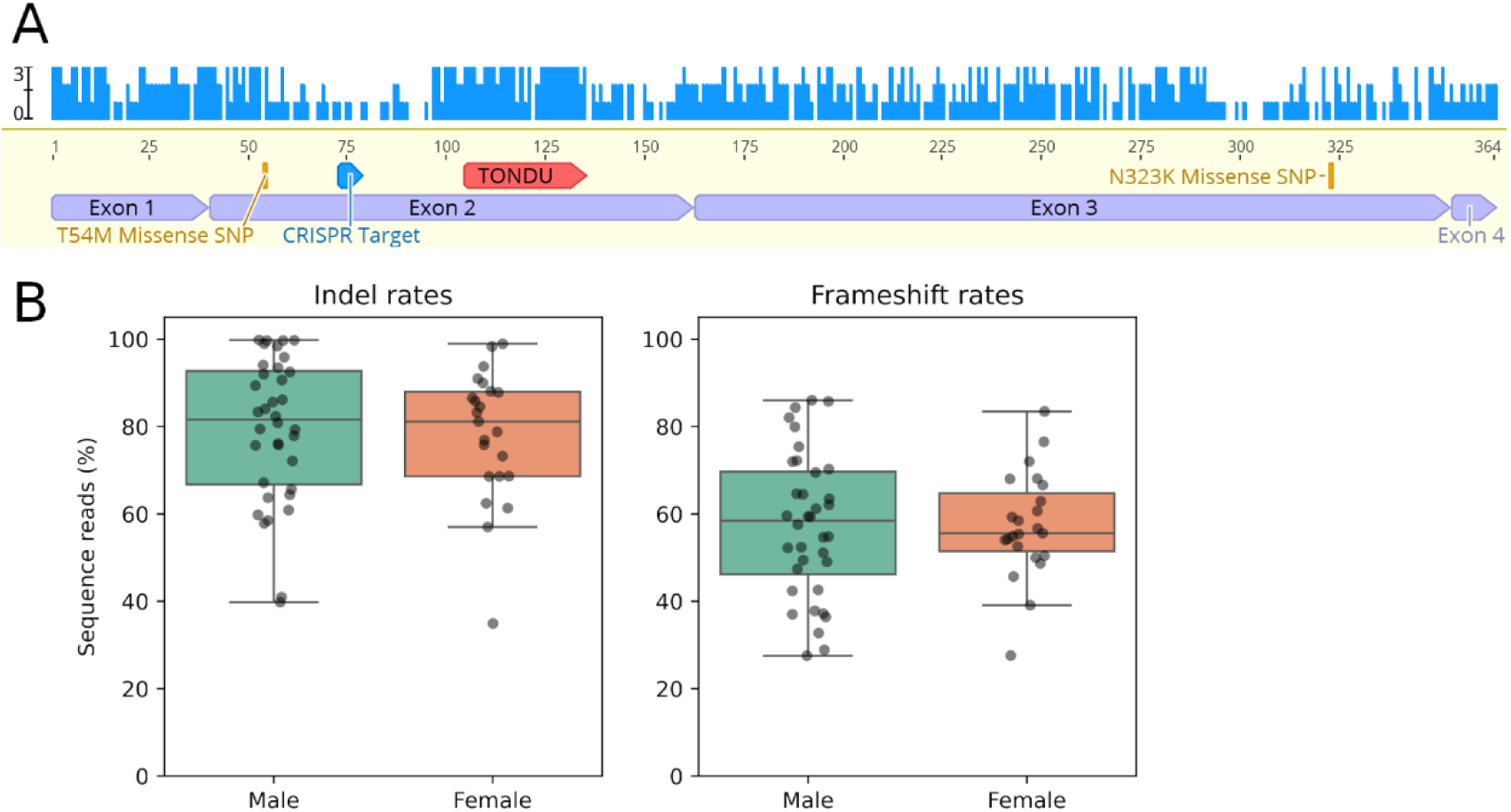
A) Annotation of Vgll3a protein sequence, indicating the locations of two missense mutations associated with pubertal timing (yellow), a TONDU domain (orange), CRISPR target site (blue) and exon locations. Blue bars at the top show the number of amino acid matches (sequence identity) per position between Atlantic salmon and Northern Pike, Zebrafish and Human. **B)** Showing total indel rates at the CRISPR target site for male and female crispants (left) and corresponding frameshift rates (right).

### 2.2. Male maturation

#### 2.2.1. F0 (2016) crispant males

A freshwater post-smolt maturation regime was applied in February 2018 (Figure 2). Males were sampled two weeks after regime onset and again in March, and September (Figure 3). Pituitary *fshb* expression and plasma 11-KT were significantly lower in crispant males compared to control males early after regime onset, while GSI did not differ. By September 2018, crispants displayed significantly fewer precociously maturing males than controls, coincident with lower 11-KT (Figure 3).

**Figure 2.**
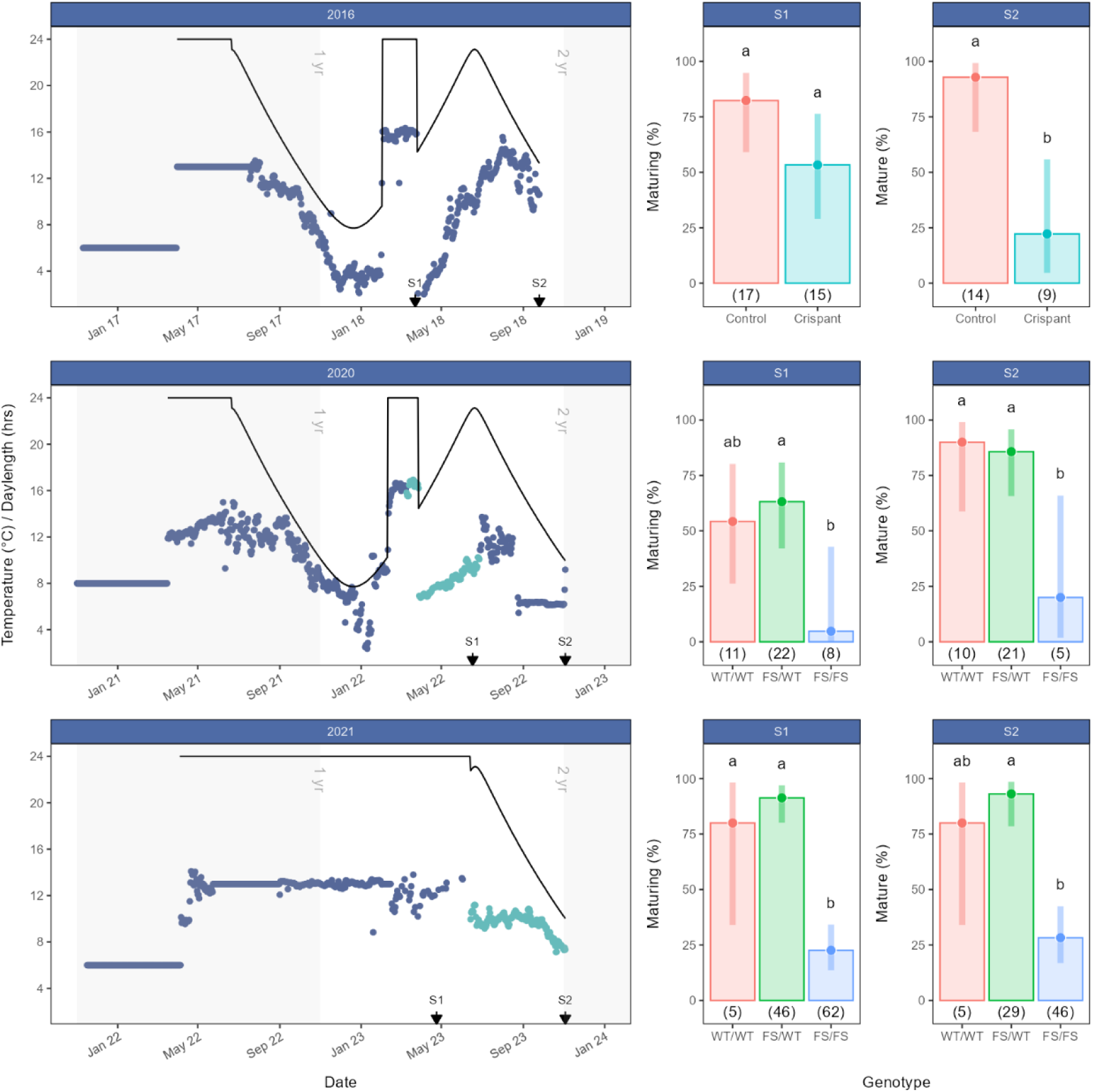
Pubertal development and final maturation observed in salmon males (1,5-2-year-old) in *vgll3a* crispants (top) and in *vgll3a* F1 fish for year-classes 2020 (middle) and 2021 (bottom) exposed to early maturation regimes. Probability of entering male maturation (Maturing, S1) and probability of mature males (S2) are shown as the estimated marginal means (+/- 95% confidence interval). Red, cyan, green and blue bars indicate *vgll3a*^+/+^ (WT/WT), crispants, *vgll3a*^+/-^ (FS/WT) and *vgll3a*^-/-^ (FS/FS), respectively. Number of individuals are presented in parentheses below each bar. Letters (a and b) indicate significant differences (P<0.05). Blue and cyan dots indicate water temperature in freshwater and brackish water (25 ppt), respectively. Arrows indicate sampling timepoints (S1 and S2).

**Figure 3.**
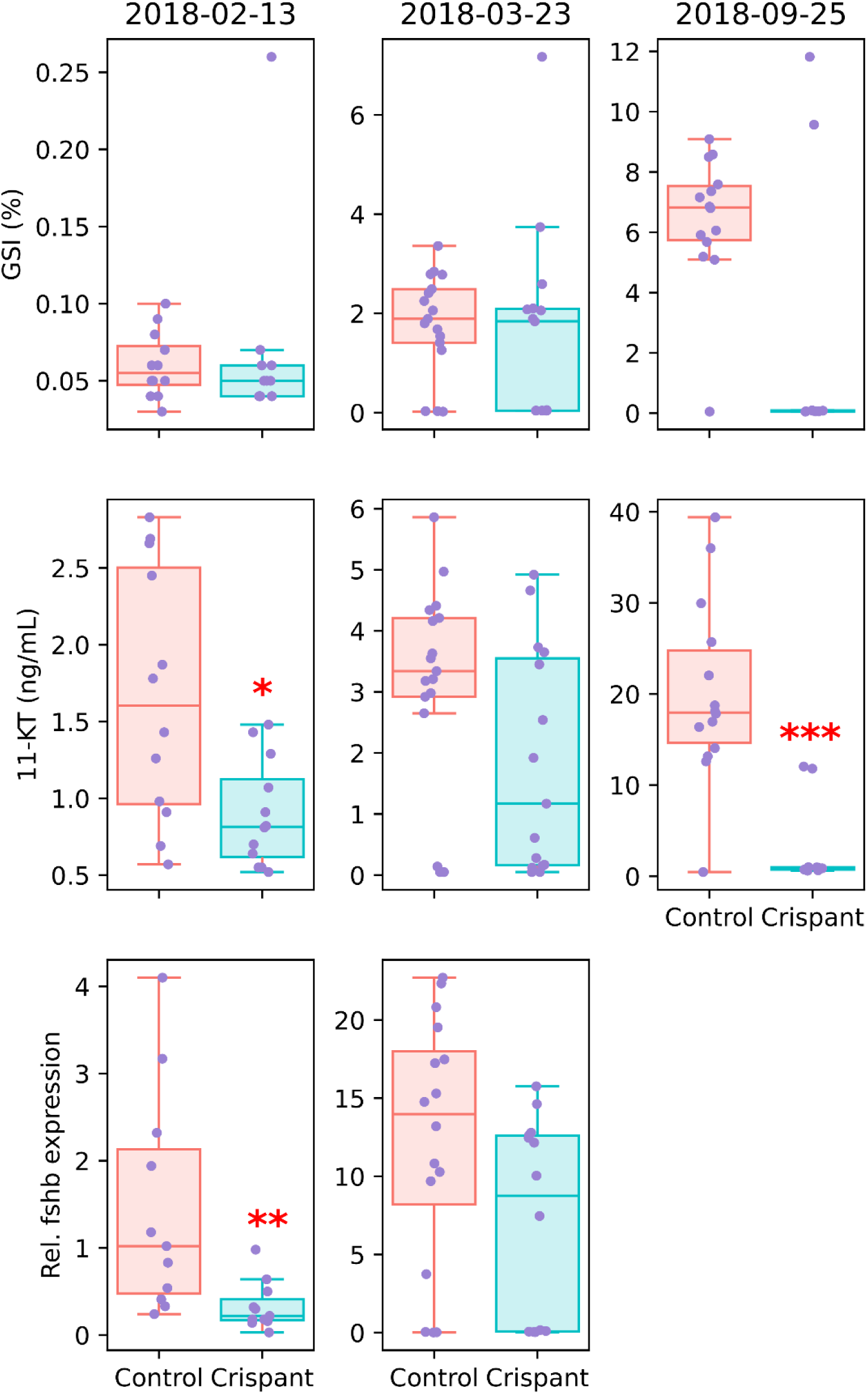
GSI, 11-KT and pituitary expression of *fshb* in controls and *vgll3a* crispant males at three timepoints. * = P < 0.05, ** = P < 0.01, *** = P < 0.001.

#### 2.2.2. F1 (2020YC and 2021YC) males

Males obtained from the 2020 year-class were monitored for maturation at two time points after ending the maturation regime in the spring of 2022; in June (S1) for pubertal assessment and in November (S2) for spawning assessment. At both timepoints we found a significantly higher probability of maturation in v*gll3a^+/+^* and *vgll3a^+/-^*as opposed to *vgll3a^-/-^* males (see Figure 2).

In the second F1 generation (2021 year-class), in two-year-olds, pubertal status was assessed in spring (S1) and at spawning in the fall (S2) revealing a significantly lower maturation rate in the v*gll3a^-/-^* group compared to *vgll3a^+/-^* (see Figure 2). We also observed a lower probability of maturation in the v*gll3a^-/-^* group compared to WT. This was only significant at the spring sampling, probably due to the low number of WT fish in the experiment. In spring, the levels of 11-KT in the plasma of v*gll3a^-/-^* (FS/FS, Figure 4A) were significantly lower compared to *vgll3a^E/-^* and *vgll3a^L/-^* males. This was also reflected in the maturation as assessed by ultrasound, in which v*gll3a^-/-^*displayed significantly lower maturation rates compared to *vgll3a^E/-^* and *vgll3a^L/-^* males (Figure 4B), clearly correlating with 11-KT concentrations.

**Figure 4.**
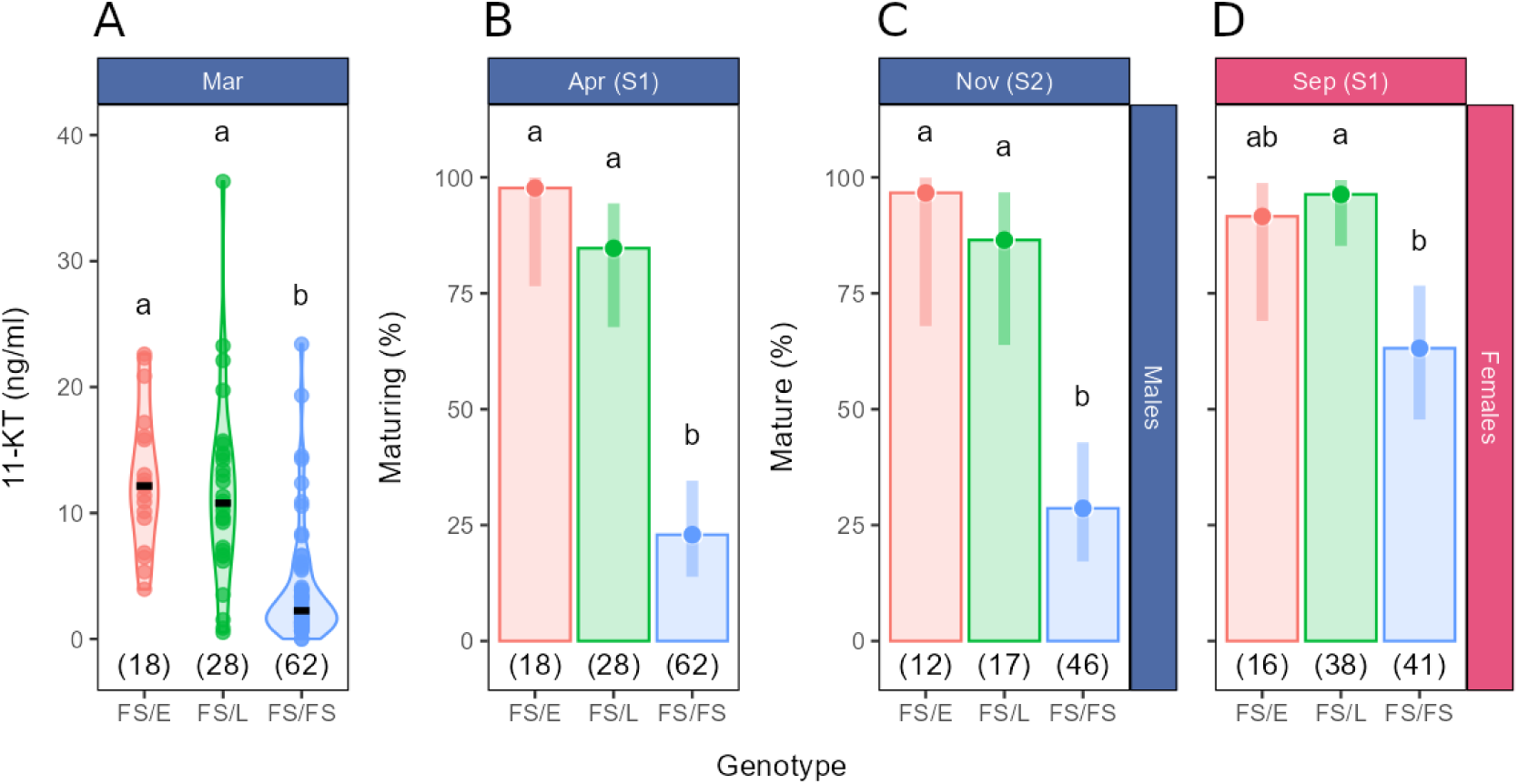
In the 2021 year-class, FS/WT (*vgll3a^+/-^*) groups were split into FS/E (*vgll3a^E/-^*) and FS/L (*vgll3a^L/-^*) depending on presence of the E or L allele in addition to a single FS allele. **A)** Concentration of 11-KT in male plasma in March 2023. **B)** Probability of entering male maturation in April 2023 (S1) and **C)** probability of mature males in November 2023 (S2), see Figure 2 for details on S1 and S2. **D)** Probability for females to mature at spawning time (October-November 2024, S1), see Figure 5 for details on S1. Probabilities (B-D) are shown as the estimated marginal means (+/- 95% confidence interval).

### 2.3. Female maturation

Assessment of maturation in four-year-old female crispants (Figure 5) showed that significantly fewer crispants matured compared to controls. In three-year-old females from the 2020 year-class, the *vgll3a^-/-^* (FS/FS) group displayed less maturation in comparison to the *vgll3a^+/+^* (WT/WT) group. In the 2021 year-class, the probability of maturation in *vgll3a^-/-^* (FS/FS) females was reduced in comparison to *vgll3a^+/-^* (FS/WT) females (Figure 5). Furthermore, a sufficiently high number of individuals was available from this year-class to also assess the allele specific contribution to maturation. Compared to the *vgll3a^L/-^* group, the *vgll3a^-/-^* females displayed significantly lower maturation rates. No difference in maturation rate was observed between *vgll3a^L/-^* and *vgll3a^E/-^*females (Figure 4D).

**Figure 5.**
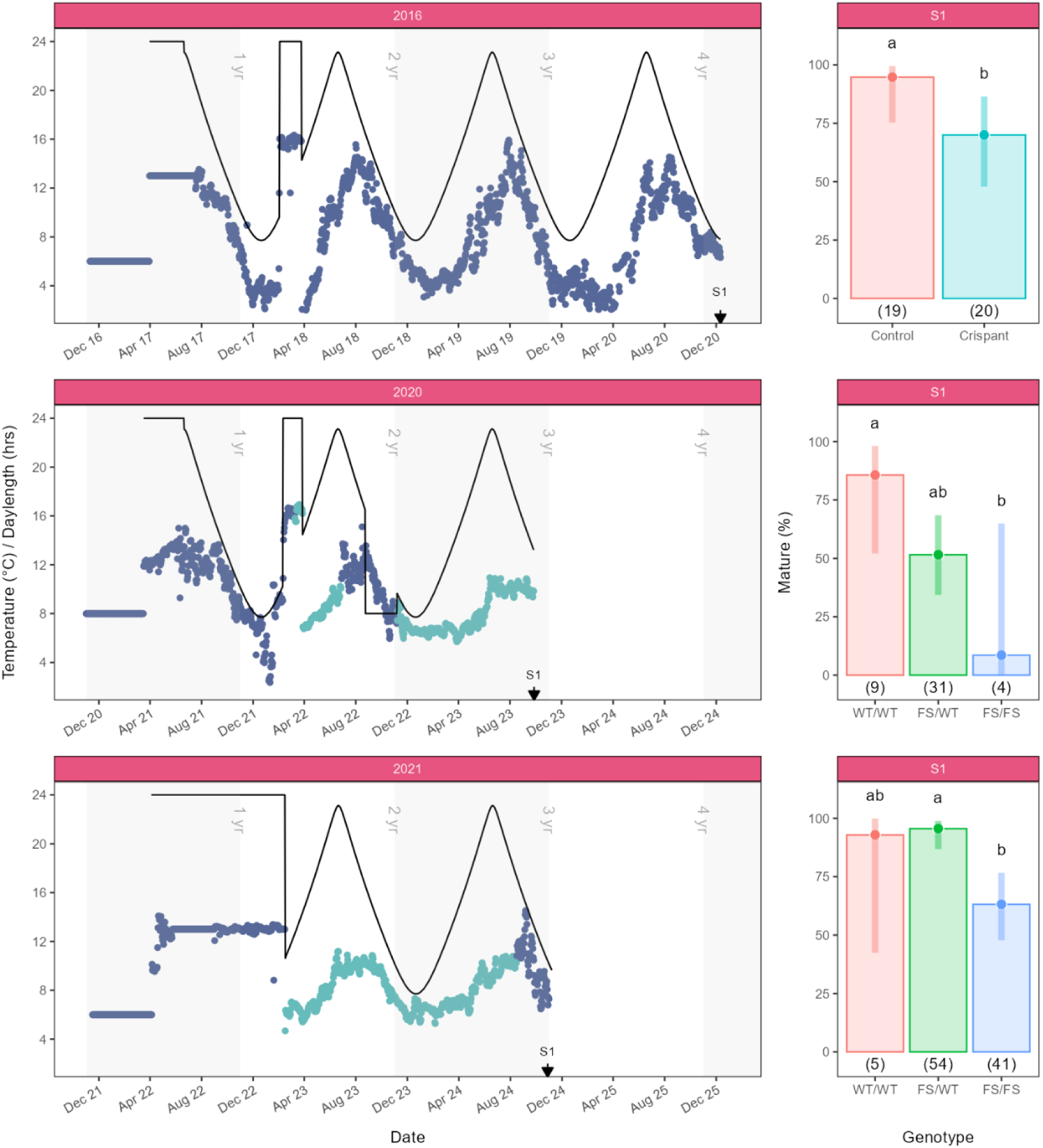
Female maturation was assessed lethally for 2016 YC (top) and 2021 YC (bottom) or by ultrasound for 2020 YC (middle) at spawning time at 3 and 4 years of age. In both 2020 and 2021 cohorts, three-year-old *vgll3a^-/-^* females showed less maturation compared to either *vgll3a*^+/+^and *vgll3a^+/-^*. In four-year-old females from 2016 YC, significantly fewer crispants matured versus controls. Probability for females to mature is shown as the estimated marginal means (+/- 95% confidence interval). Red, cyan, green and blue bars indicate *vgll3a*^+/+^ (WT/WT), crispants, *vgll3a*^+/-^ (FS/WT) and *vgll3a*^-/-^ (FS/FS), respectively. Number of individuals are presented in parentheses below each bar. Letters (a and b) indicate significant differences (P<0.05). Blue and cyan dots indicate water temperature in freshwater and brackish water (25 ppt), respectively. Arrows indicate sampling timepoints (S1).

## 3. Discussion

Our results demonstrate that disrupting the *vgll3a* gene consistently reduced the rates of sexual maturation in both male and female Atlantic salmon. This effect was evident across multiple year classes and under different maturation stimulation regimes. In both sexes, the complete loss of *vgll3a* function was required to reduce rates of maturation, while heterozygous individuals matured at similar rates as controls. These results confirm that *vgll3a* is important for activating the reproductive axis and strongly supports its role as a positive regulator of pubertal onset in salmon. Previous genome-wide association studies identified *vgll3a* as the major locus explaining variation in age at maturity (1,2) but the function in maturation has remained unclear.

This study fills part of this knowledge gap by providing direct functional evidence that *vgll3a* promotes sexual maturation in Atlantic salmon. This finding is both consistent with and divergent from recent observations in killifish (22). In killifish, mutations in exon 1 that introduced a premature stop codon increased the incidence of precocious male maturation. The authors hypothesized that this phenotype resulted from an increased expression of a shorter *vgll3* isoform. Similarly, a previous study in Atlantic salmon reported the existence of a comparable short *vgll3a* isoform (26). However, in contrast to our findings, that study proposed that the expression of the short *vgll3a* isoform is associated with the L allele, thereby maintaining the gonad in an immature state. We examined RNA-seq data from 10 immature testis samples from fish having EL genotypes for evidence of allele-dependent differences in isoform expression. In contrast to the previous salmon study, we found that the short isoform exhibited similar expression levels of the L and E alleles (Supplementary File S4). Thus, our data does not support a role for the short isoform in timing of maturation in Atlantic salmon. Interestingly, introduction of a nonsense mutation in exon 3 of killifish *vgll3* marginally (P=0.049, n=6) inhibited male maturation. This phenotype is consistent with our observations in Atlantic salmon, although the effect appears less pronounced in killifish. Notably, and in contrast to our findings, heterozygous exon 3 mutant killifish exhibited an increase in precocious maturation, a phenotype that was not observed in Atlantic salmon. In our study, heterozygous males (and similarly females) displayed maturation rates comparable to those of the controls, indicating that loss of both functional *vgll3a* alleles is required to reduce maturation in salmon, whereas partial loss of function appears sufficient to alter maturation timing in killifish. By generating functional knock-outs, our study provides direct experimental evidence that complete loss of *vgll3a* function reduces the incidence of early maturation, demonstrating that Vgll3a actively promotes the onset of puberty in Atlantic salmon males, in line with what has been observed for the vgll3 exon 3 mutant killifish.

In the 2021 year-class, we had an even distribution of E and L alleles, allowing comparison between *vgll3a^E/-^* and *vgll3a^L/-^*heterozygotes. We found no maturational differences between these genotypes; both matured at similar rates. These results continue to challenge (10,27) the interpretation of sex-dependent dominance at *vgll3a* proposed by Barson et al. (2015) (1). Consistent with our results, several independent studies across wild and domesticated Atlantic salmon populations have failed to detect a stable or generalizable sex-dependent dominance pattern at *vgll3a* E and L alleles, reporting instead additive effects, weak or absent dominance, or environment- and strain-specific genotype–phenotype relationships (10,28–31). In our functional experiments, disruption of *vgll3a* altered the probability of maturation in both males and females, while heterozygotes matured at normal rates, indicating that *vgll3a* function is not dose-dependent. This aligns with previous findings that dominance relationships at *vgll3a* are weak, variable, or absent under certain environmental or genetic backgrounds (10,29). Moreover, the inherent sexual dimorphism in maturation timing was maintained, with males maturing earlier than females even in the absence of *vgll3a* function. Together, these results suggest that the sex-dependent dominance pattern inferred from population-level association studies does not arise from intrinsic allelic dosage effects or sex-specific *vgll3a* function.

Reduced *fshb* expression in the pituitary of *vgll3a* crispant males points to *vgll3a*’s involvement in regulating the BPG axis. Fsh is an important regulator of male maturation in salmon (17), and earlier studies have shown similar *vgll3a* genotype-dependent differences in *fshb* expression (15,16). This data supports the notion that Vgll3a modulates maturation by stimulating *fshb* expression in the pituitary. Fsh stimulates testicular androgen production and elevates 11-KT plasma levels in different fish species (e.g. salmon (32), zebrafish (33) and sea bass (34)). We consider the reduced 11-KT plasma levels found in males after the complete loss of Vgll3a as reflecting a down-stream effect of reduced Fsh stimulation in these mutants. However, since salmon still can mature, albeit at a reduced rate, also in the absence of Vgll3a, we conclude that Vgll3a is an important, but not an essential or required, activator of the maturation of the BPG axis. Our earlier studies reported a decrease of testicular *vgll3a* expression occurring with the onset of puberty in salmon (13). This previous observation may reflect a dilution of the testicular *vgll3a* mRNA signal, produced by the somatic Sertoli cells being increasingly outnumbered by the germ cells produced during spermatogenesis. Alternatively, lower testicular *vgll3a* levels after the onset of puberty may indicate that *vgll3a* is down-regulated after having contributed to triggering the initiation of testis maturation. Hence, the clear reduction in maturation frequency among *vgll3a^-/-^* fish, together with the endocrine data, supports the view that one of the *vgll3a* functions is to regulate the BPG axis through stimulation of gonadotropin production and steroidogenesis. Multi-omics analyses show that *vgll3a* genotypes influence expression of a steroidogenic enzyme, *cyp17a1* (35), which is critical for androgen production. Hence, Vgll3a likely acts upstream of this enzyme, coordinating its expression through Hippo/YAP/TEAD transcriptional control, possibly via elevated levels of *fshb* in the pituitary. Expression of *vgll3a* in Sertoli cells of the testis may modulate the release of a yet unidentified factor(s) into the plasma, which in turn affects pituitary expression of *fshb*. This mechanism could be related to recent findings of cholecystokinin (Cck)-induced *fshb* expression in, and Fsh release from, the pituitary (36,37). However, these so far hypothetical links between testicular Vgll3a and pituitary function remain to be substantiated in future studies.

Female *vgll3*a^-/-^ knockouts showed lower frequency of maturation, consistent with the male results. In three-year-old F1 females and four-year-old crispant females, we observed strong genotype-dependent differences in maturation rates, with crispants and *vgll3a^-/-^* fish showing less maturation. Heterozygotes (*vgll3*a^+/-^) matured normally, reinforcing that both alleles must be lost to reduce maturation frequencies.

VGLL3 acts as a cofactor for TEAD transcription factors within the Hippo signaling pathway, a conserved system regulating organ growth and cell fate (7). In salmon, *vgll3a* expression correlates with Hippo pathway genes including *yap1* and *tead3* (13). This mechanism aligns with the heterochronic model proposed by Verta et al. (2024) (35), where the *vgll3a* genotype affects the timing of activation of testicular developmental networks. Our results therefore provide functional evidence that *vgll3a* acts by contributing to the developmental activation of puberty, controlling Hippo pathway activity, which may increase *fshb* in the pituitary (15,16) and result in steroidogenic gene expression (38). From an aquaculture perspective, a lower incidence of precocious maturation is favorable since maturation causes reduced growth, lower flesh quality, and higher mortality (17). The loss of Vgll3a may therefore be advantageous, for example in RAS facilities, where precocious maturation commonly results in welfare problems (23). However, before considering practical applications, a careful assessment of potential, pleiotropic effects of Vgll3a will be required. After all, previous work suggested that Vgll3a may influence growth, metabolic regulation, or immune function (18–20). Moreover, the long-term fertility of *vgll3a^-/-^* fish remains to be studied. So far, it seems that the *vgll3a* knockout model may offer a promising framework for precision breeding aimed at improving welfare and production stability.

Although our study demonstrates clear causal effects of *vgll3a* on maturation, the molecular mechanisms connecting Vgll3a function to pubertal maturation remain partly unresolved. Which tissue is the key source of Vgll3a-mediated effects on maturation? Previous results from our group and from others point to all levels of the BPG axis. Expression of *vgll3a* is regulated within the testis, and particularly within the Sertoli cells upon entry into puberty, but the gene is also highly expressed in the heart and gill. The latter two organs are key for delivering oxygen to the body, another pathway linked to size at maturation (39). Because the paralog *vgll3b* was not targeted, potential compensatory effects cannot be ruled out. In addition, endocrine signals are necessary to transmit reproductive cues, but the Vgll3a-triggered factor(s) mediating the potential feedback between the testis, brain, and pituitary remain(s) to be established. Overall, at this point, it seems possible that a multi-tissue endocrine/metabolic network may be involved in mediating Vgll3a effects on maturation, a possibility requiring significant future research efforts.

Genetic variation at the human VGLL3 locus has been associated with the timing of puberty in females (21); however, as discussed above, the mechanistic role of VGLL3 in this process remains unclear. Based on the results obtained in this study, salmon may provide a useful comparative model to investigate this function, as VGLL3 has been implicated in multiple biological processes in humans. In particular, it has been linked to female-specific autoimmune pathways (20) and to cancer biology, where it regulates cellular proliferation and cell-state transitions across several tissue types (19). Following genome duplication in salmonids, *vgll3* paralogs may have undergone sub-functionalization, whereby *vgll3a* retained a role in the regulation of maturation, while the alternative paralog, *vgll3b*, may contribute to other pleiotropic functions. This is further supported by previous studies showing that only genetic variation in *vgll3a* is associated with age at maturity (2) and that only this paralog is regulated in the gonad upon puberty initiation (13).

## 4. Conclusion

Loss of *vgll3a* function reduces the predisposition to enter early maturation in Atlantic salmon. In both sexes, complete knockout of *vgll3a* (*vgll3*a^-/-^) is necessary to reduce the frequency of maturation, while heterozygous (*vgll3a*^+/-^) individuals mature at normal rates. Thus, the effect of *vgll3a* disruption is identical in males and females: only homozygous knockouts display a reduced incidence of maturation. This finding provides functional evidence that Vgll3a acts as one of the important activators of sexual maturation in salmon.

Our results also show that *vgll3a* function is not dose-dependent in our experimental conditions, as heterozygous knockouts (*vgll3a*^+/-^) mature at normal rates, independent of the remaining functional allele being the E or L variant. Moreover, the characteristic sex difference in age at first maturity remains unchanged, males still mature earlier (typically at 1–2 years) than females (3–4 years), indicating that *vgll3a* affects the overall probability or readiness to mature rather than the relative timing between sexes. At a broader scale, these results suggest that Vgll3 may play a role in promoting puberty in vertebrates and that salmon may serve as a useful model for investigating this function, potentially due to the existence of two *vgll3* paralogs in salmon.

## 5. Materials and methods

### 5.1. Guide RNA design and synthesis

Cas9 mRNA and Guide RNA (gRNA) for use with CRISPR/Cas9 was synthesized as described by Edvardsen et al. 2014 (40). The *vgll3a* gRNA (5’-GAGGCTGGCTGGGGAGAAGG-3’) was designed to target exon 2 of *vgll3a* (GeneID:106586514) on chr 25 (Figure 1), while avoiding homology with the *vgll3b* paralog on chr 21. gRNA targeting *slc45a2* was used for co-injection with *vgll3a* guide to produce albino phenotype for screening, as previously described (40,41).

### 5.2. Experimental animals and rearing conditions

An overview over all experiments in terms of dates, temperatures, light conditions and resulting rates of maturation, can be seen in Figure 2 (males) and Figure 5 (females). At the samplings where ultrasound was used to estimate gonad size the fish were anesthetized with 100 mg/L Finquel vet. (tricaine methane sulfonate). Prior to all end-point samplings, clove oil (1 mL/100 L Aqui-S vet.) was applied as a sedative and the fish were anesthetized with 100 mg/L Finquel vet. At sampling, length and weight was recorded and blood was taken before the fish were sacrificed by severing the medulla oblata. Pituitaries and gonads were excised and the gonadosomatic index (GSI, % of body weight that is gonad) was calculated. Biometrics obtained from the experiments are found in Supplementary File S1.

#### 5.2.1. 2016 year-class (F0 crispants)

Eggs and sperm were provided by Aquagen. The temperatures and photoperiods experienced by the males and females are found in Figure 2 and 5, respectively. Briefly, on the 10th of November 2016 the eggs were fertilized and microinjected with 50 ng/µl *vgll3a* gRNA, 50 ng/µl *slc45a2* gRNA and 150 ng/µl Cas9 mRNA, as previously described (40). Injected eggs were incubated in the hatchery at 6 °C. The fish were moved to ∼13 °C and LD 24:0 for start feeding on the 31^st^ of March 2017. The photoperiod was set to simulated natural photoperiod (SNP, based on civil daylength, 62 °N) on the 21^st^ of June 2017 and the temperature was switched to ambient on the 3^rd^ of August 2017. Between the 31^st^ of January and 2^nd^ of February 2018 the temperature was increased from ambient (∼4 °C) to 16 °C and the photoperiod was switched from SNP to LD 24:0 to stimulate post-smolt maturation which is strongly regulated by *vgll3a* genotypes for late and early maturation (11). The fish remained in these conditions until the 26^th^ of March 2018 when they were moved back to ambient temperature (∼2 °C) and natural light. The fish were kept on freshwater (0.1 ppt) throughout and fed 20% excess based on their body weight and expected growth rates. The fish were sampled on 3 occasions in 2018, on the 13^th^ of February, the 23^rd^ of March and the 25^th^ of September. Lethal sampling was performed on each date. Blood was collected and the fish were sacrificed by cutting the medulla oblongata. Blood samples were centrifuged at 14,000 x *g* at 4 °C for 2 min, plasma was collected and stored in −80 °C until further analyses. Pituitaries were stored in RNA later (Thermo Fisher Scientific) at 4 °C or −80 °C until further use. The length, weight and gonad weight of all sampled fish were recorded. The remaining fish were further held in fresh water, at ambient temperature and photoperiod for two years until the females reached maturation autumn 2020. Only a few males were kept for cross out (See section 5.2.3 and 5.2.4). On the 11^th^ of December 2020 all remaining fish (see Supplementary File 1) were terminated, and maturation was recorded.

#### 5.2.2. 2017 year-class (F0 crispants)

Eggs and sperm were provided by Aquagen. On the 2^nd^ and 3^rd^ of November 2017 eggs were fertilized and microinjections were performed as described for the 2016 year-class. Fish showing full pigmentation without albino spots on the skin were removed, as they were assumed to have low or zero mutation rates. The fish were then reared at the same regimes as year-class 2016. These fish were only used for breeding the F1 generations, and no specific samplings were done on them, except for implanting passive integrated transponder (PIT) tags and fin clipping for genotyping in one-year-old smolts. In summary, fish were exposed to a standard salmon regime: in freshwater at 6 °C until start-feeding, followed by 13 °C and continuous light for three months, and then natural light and temperature until smoltification at 1.5 years of age, followed by seawater transfer and rearing under natural light and temperature until maturity.

#### 5.2.3. F1 (2020 year-class)

Eggs from two female crispants from the 2016 year-class and sperm from two male crispants from the 2017 year-class (see Supplementary File S2) were used to produce four families in the F1 generation. All four parents were selected based on presence of frameshift (FS) mutations (68-79% FS rate) in fin-clip DNA. Genotyping of fish from this mix of families enabled selection of *vgll3a^+/+^* (WT/WT), *vgll3a^+/-^* (FS/WT) and *vgll3a^-/-^* (FS/FS) that could be used in maturation experiments. An overview of the experimental fish groups can be found in Supplementary File S1. The temperatures and photoperiods experienced by the males and females are shown in Figure 2 and 5, respectively. Briefly, the eggs were fertilized on the 27^th^ of November 2020 and incubated at 8 °C. The fish were moved to ∼12 °C and LD 24:0 for start feeding on the 17^th^ of March 2021. The temperature was switched to ambient (∼12 °C) on the 4^th^ of June 2021 and the photoperiod set to SNP on the 21^st^ of June 2021. On the 17^th^ of January 2022 the fish were moved from ambient (∼4 °C) to ∼10 °C, before being moved up to 16 °C, between the 10^th^ and 16^th^ of February 2022. The photoperiod was also switched from SNP to LD24:0 on the 10^th^ of February 2022 to stimulate post-smolt maturation. On the 10^th^ of March 2022, the salinity was changed from freshwater to brackish (25 ppt) water. The fish remained on 16 °C brackish water and LD 24:0 until the 28^th^ of March 2022 when they were moved to ∼7 °C brackish water and SNP. Due to the relatively low number of males, we could not perform lethal samplings during the maturation inducing regime. However, on the 17^th^ of June 2022, we performed an ultrasound evaluation of maturity in all males. The salinity was changed from brackish to freshwater on the 28^th^ of June 2022. On the 16^th^ of August 2022, the males and females were split. The females remained on ambient freshwater, but the photoperiod was changed to LD 8:16 on the 24^th^ of August 2022, before they went back on ∼8 °C brackish water and SNP on the 7^th^ of November 2022. Due to an incident with a waterpipe at the end of August 2022, where we unfortunately lost 2, 1 and 3 v*gll3a^+/+^*, *vgll3a^+/-^*and *vgll3a^-/-^* individuals, respectively, fewer fish were monitored at the last timepoint as opposed to the June maturation monitoring. The males were moved to ∼6 °C freshwater and SNP on the 23^rd^ of August 2022 where they stayed at ambient temperature until their final sampling on the 2^nd^ of November 2022, where maturity status was assessed by gently squeezing the belly to determine whether they released milt or not, in combination with visual inspection of gonads. Females were assessed for maturity on the 27^th^ of September 2023 using ultrasound.

#### 5.2.4. F1 (2021 year-class)

In the autumn of 2021, eggs and sperm from two females and two males from the 2017 year-class (see Supplementary File S2) were used to produce a second F1 generation consisting of four families. All four parents were selected based on presence of frameshift (FS) mutations (81-86% FS rate) in fin-clip DNA. An overview of the experimental fish groups can be found in Supplementary File S1. The temperatures and photoperiods experienced by the males and females can be seen in Figure 2 and 5, respectively. Briefly, the eggs were fertilized on the 16^th^ of November 2021 and incubated at 6 °C. The fish were switched to ∼13 °C and LD 24:0 for start feeding on the 6^th^ of April 2022. They remained on the same conditions until the 3^rd^ of February 2023, when the males and females were moved to separate tanks. On the 15^th^ of February 2023, the females were transferred to brackish water and SNP. On the 22^nd^ of March 2023, plasma was sampled from males for analysis of 11-KT concentrations. On the 24^th^ of April 2023, length and weight were measured, and an ultrasound evaluation of gonad maturation was done for all males to determine if they were pubertal or not. The males remained on ∼13 °C and LD24:0 until the 14^th^ of June 2023 to replicate commercial “large smolt” production, which also results in large amounts of precocious maturation in males (27). On the 14^th^ of June 2023, the males were moved back to the same tanks as the females on ∼10 °C brackish water and SNP for the remainder of the study period. On the 3^rd^ of November 2023, the maturity status in males was assessed by gently squeezing the belly to determine whether they released milt or not, in combination with visual inspection of gonads. Maturity in females was assessed at spawning time, with lethal samplings on the 28^th^ of October and 7^th^ of November 2024.

### 5.3. Genetic analyses

#### 5.3.1. Fin-clipping and DNA extraction

All fish used in this study were implanted with a PIT tag while sedated with Finquel vet. (0.1 g/L), immediately followed by removal of a small piece of the adipose fin for genetic analyses. DNA was extracted from fin tissue with Qiagen Blood and Tissue Kit (Qiagen) for the 2016 and 2020 year-classes and Biomek i5 Automated Workstation (Beckman Coulter) for the 2021 year-class.

#### 5.3.2. Mutation analysis

To assess the mutation types of *vgll3a* in crispants and mutation genotype frequencies of individuals in the F1 generations, amplicon sequencing was performed using the MiSeq (Illumina) sequencing platform. Nested PCR using Q5 High-Fidelity DNA Polymerase (NEB) was used to amplify a 350 bp genomic region including the *vgll3a* CRISPR target site (primer sequences listed in Supplementary File S3), followed by barcoding with sample-specific dual-index adapters. Samples were pooled in equal volumes, followed by gel extraction using a QIAquick Gel Extraction Kit (Qiagen). Pooled DNA was quantified using Qubit (Thermo Fisher) and adjusted to 10 nM. DNA pools were sequenced using MiSeq Kit v.3 (Illumina) with 300 bp paired-end reads. Processing of the sequenced reads was performed using custom scripts. The 5’ end of each read was required to match the target site-specific primer sequence. The reads of each read-pair were assembled to improve sequence quality, followed by mapping to the reference amplicon sequence using Muscle (v. 3.8.1551 (42)). Reads were screened for the presence of insertions and deletions (indels) and further classified into the following categories based on the effect on the reading frame: frameshift (FS or -), in-frame (IF) or no indel (WT or +). For the F1 generations, genotypes were called as follows: If an individual had reads representing a given allele with frequency > 90%, that individual was considered homozygous for that allele. If an individual had two alleles present at frequencies > 40%, that individual was considered heterozygous for those two alleles. Further, individuals were categorized as FS/FS (homozygous for FS alleles, *vgll3a*^-/-^) if having only FS alleles, and FS/WT (*vgll3a*^+/-^) if having one FS allele and one WT allele. Individuals without FS (*vgll3a^-^*) or IF (*vgll3a^IF^*) alleles were categorized as WT/WT (*vgll3a^+/+^*). Individuals having other allele combinations (*vgll3a^-/IF^*, *vgll3a^IF/IF^*and *vgll3a^+/IF^*), or not passing the filtering thresholds, were removed from the experiments and are not included in the analysis. For the 2021 year-class, the *vgll3a^+^* allele was further divided into *vgll3a^E^* (early) and *vgll3a^L^*(late) as these alleles previously have been associated with age at maturation (1,2), resulting in the genotype groups *vgll3a*^-/-^,*vgll3a^E/-^*, *vgll3a^L/-^* and *vgll3a^+/+^* (due to low numbers of WT individuals having EE, EL or LL genotypes without any CRISPR-generated mutations, such individuals were merged into a single group *vgll3a*^+/+^).

In some of the individuals in the 2021 year-class a large deletion of 442 bp was detected that removed the forward primer binding site for the sequencing primers. To correct the genotypes of individuals having this deletion, a PCR assay was designed to screen for this deletion in the 2021 year-class. The PCR reaction contained 3 primers, with two primers flanking the deletion, and a single primer overlapping the junction. The PCR was performed using GoTaq DNA Polymerase (Promega) with 0.25 µM of each primer (primers are listed in Supplementary File S3), followed by gel electrophoresis to inspect PCR-product sizes. Presence of a large band (753 bp) indicated WT allele, while presence of a medium sized band (311 bp) indicated a large deletion. The additional presence of a small band (175 bp) indicated the presence of the specific deletion of 442 bp. The large deletion was considered as a *vgll3a^-^* (FS) allele. Sequencing-based genotypes were corrected for all individuals positive for the large deletion.

#### 5.3.3. Allelic discrimination

Genotyping of a natural single nucleotide polymorphism causing an amino acid replacement on aa 323 in Vgll3a associated with age at maturation was performed according to Ayllon et al 2019 (10). Briefly, an allelic discrimination assay was performed using TagMan Genotyping Master Mix (Thermo Fisher Scientific) with primers and probes shown in Supplementary File S3. The assay was run on QuantStudio 5 (Applied Biosystems).

#### 5.3.4. Genetic sex

Genetic sex was determined by assaying the presence of exons 2 and 4 of the *sdy* gene in genomic DNA as described by Ayllon et al 2019 (10), but with corrected primers and probes (shown in Supplementary File S3). All fish included in this study were analyzed with this assay. Most fish were dissected when sacrificed, and presence of ovaries or testes was noted. In the few fish that showed discrepancy between genetic sex and phenotypic sex, the phenotypic sex was used in data analysis.

5.4 **Androgen plasma concentrations and pituitary *fshb* gene expression analyses** Plasma 11-ketotestosterone (11-KT) was measured by ELISA in males in crispants and in the 2021 year-class, as described previously (25). Expression of *fshb* in pituitaries of crispant and control males in the 2016 year-class was analyzed by qPCR, as described previously (38).

### 5.5 Maturation status

Maturation status was assessed in 2016YC males at two lethal samplings, with visual inspection of gonads in combination with GSI and 11-KT plasma levels. In 2020YC and 2021YC, male maturation status was assessed by ultrasound imaging (Mindray E7, Adcare) in the spring or summer for detecting the presence of visible maturing gonads, and in the fall by lethal sampling and visual inspection of the gonads (43). Females of the 2016 and 2021 year-classes were dissected at spawning time and visually scored as mature or not. In the 2020 year-class, females were scored as mature or not based on presence of mature gonads identified by ultrasound assessment.

### 5.6 Statistics

The maturation data was analyzed using logistical regression within each year-class, sex, and timepoint separately. It was performed using either the “glm” function from the “stats” package included as part of base R (44) or the “brglm” function from the “brglm” package (45), with a binary outcome and a probit link function. Bias reduction (“brglm”) was used when there was near complete separation of the data within a group leading to an inability to predict confidence intervals from estimated marginal means (EMMs). For all models, phenotype (immature vs mature) was set as the dependent variable, with genotype (2 levels for the 2016 year-class, “wild” vs “crispant” and 3 levels for the 2020 and 2021 year-classes, “WT/WT”, “FS/WT”, “FS/FS”) as the independent variable. The “emmeans” package (46) was used to calculate EMMs from each model with the intervals back transformed from the probit scale. For the year-classes with more than 2 genotypes, the “cld” function from the “multcomp” package (47) was used to undertake contrasts on the probit scale with a tukey correction for comparing three estimates. The same approach was used to assess *vgll3a* allele effects in the 2021 year-class. Statistical analyses of 11-KT, GSI and *fshb* gene expression were performed using Mann-Whitney U tests (two-sided) from the scipy.stats module in Python.

## Author contributions

EK, KS, FA, AW, and RBE performed the CRISPR experiments. EK, TF, SB, AW, and PGF performed the maturation experiments. EK, PV, SB, EA, and BN carried out the molecular and hormone analyses. EK, TF, EA, RWS, and AW analyzed the data and wrote the manuscript. All authors reviewed and approved the final version of the manuscript.

## Acknowledgements

We thank Audun Ø. Pedersen, Anne Hege Straume, Anne Torsvik, Sara Olausson, Ivar Helge Matre, Tone Knappskog, Christine Sørfonn and Lise Dyrhovden for help with fish rearing, samplings and labwork. This work is funded by the Norwegian Research Council (project numbers 226221, 254783 and 324890).

## Ethics Approval

The use of the experimental animals in this study was performed in accordance with the Norwegian Animal Welfare Act of June 19, 2009, in force from January 1, 2010. All the fish reared were approved by the Norwegian Food Safety Authority (https://www.mattilsynet.no, permit number 29251, 22082 and 5741).

