## Supplementary File S1 for "Vgll3a promotes sexual maturation in male and female Atlantic salmon"

**Biometrics and genotypes for 2016 year-class**

| PIT tag | Group | Sex | WT (%) | In-frame (%) | Frame-shift (%) | Total read count | Sampling date | Length (cm) | Weight (g) | GSI (%) | 11-KT (ng/mL) | *fshb* (rel. expres-sion) | Maturation status |
|---|---|---|---|---|---|---|---|---|---|---|---|---|---|
| 900119000341511 | Control | Male | | | | | 2018-02-13 | 24 | 160 | 0.05 | 1.43 | 1.18 | Immature |
| 900119000341809 | Control | Male | | | | | 2018-02-13 | 23.2 | 151 | 0.09 | 1.87 | | Immature |
| 900119000341411 | Control | Male | | | | | 2018-02-13 | 20.3 | 99 | 0.06 | 1.78 | 1.02 | Immature |
| 900119000341651 | Control | Male | | | | | 2018-02-13 | 23.5 | 153 | 0.05 | 0.57 | 0.33 | Immature |
| 900119000341359 | Control | Male | | | | | 2018-02-13 | 22.5 | 129 | 0.03 | 2.83 | 0.24 | Immature |
| 900119000341822 | Control | Male | | | | | 2018-02-13 | 24.6 | 183 | 0.07 | 0.69 | 2.32 | Immature |
| 900119000341898 | Control | Male | | | | | 2018-02-13 | 22 | 124 | 0.06 | 2.66 | 0.41 | Immature |
| 900119000341388 | Control | Male | | | | | 2018-02-13 | 21.4 | 109 | 0.05 | 0.91 | 0.54 | Immature |
| 900119000341258 | Control | Male | | | | | 2018-02-13 | 24.8 | 199 | 0.1 | 2.45 | 3.17 | Immature |
| 900119000341619 | Control | Male | | | | | 2018-02-13 | 22 | 126 | 0.04 | 0.98 | 1.94 | Immature |
| 900119000341499 | Control | Male | | | | | 2018-02-13 | 22.3 | 143 | 0.08 | 2.69 | 4.1 | Immature |
| 900119000341162 | Control | Male | | | | | 2018-02-13 | 22.5 | 140 | 0.04 | 1.26 | 0.83 | Immature |
| 900119000341328 | Crispant | Male | 20.72 | 9.8 | 69.48 | 40527 | 2018-02-13 | 19 | 84 | 0.07 | 0.64 | 0.3 | Immature |
| 900119000341314 | Crispant | Male | 17.66 | 32.92 | 49.43 | 34403 | 2018-02-13 | 21.7 | 125 | 0.04 | 1.48 | 0.64 | Immature |
| 900119000341628 | Crispant | Male | 9.36 | 26.01 | 64.63 | 34724 | 2018-02-13 | 24.2 | 116 | 0.04 | 0.7 | 0.16 | Immature |
| 900119000341386 | Crispant | Male | 0.32 | 13.65 | 86.03 | 36485 | 2018-02-13 | 20.5 | 101 | 0.04 | 0.55 | 0.22 | Immature |
| 900119000341297 | Crispant | Male | 1.02 | 13.21 | 85.77 | 31643 | 2018-02-13 | 19.3 | 84 | 0.06 | 0.55 | 0.18 | Immature |
| 900119000341944 | Crispant | Male | 40.22 | 5.12 | 54.66 | 28918 | 2018-02-13 | 21 | 113 | 0.04 | 0.91 | 0.32 | Immature |
| 900119000341932 | Crispant | Male | 0.25 | 15.38 | 84.37 | 33414 | 2018-02-13 | 19.2 | 82 | 0.05 | 0.82 | 0.14 | Immature |
| 900119000341313 | Crispant | Male | 6.52 | 29 | 64.48 | 35530 | 2018-02-13 | 18.8 | 79 | 0.05 | 0.52 | 0.03 | Immature |
| 900119000341575 | Crispant | Male | 24.18 | 16.36 | 59.46 | 33303 | 2018-02-13 | 21.2 | 113 | 0.04 | 1.43 | 0.5 | Immature |
| 900119000341607 | Crispant | Male | 39.12 | 11.84 | 49.04 | 36671 | 2018-02-13 | 22 | 122 | 0.06 | 1.07 | 0.98 | Immature |
| 900119000341746 | Crispant | Male | 0.19 | 29.57 | 70.24 | 32753 | 2018-02-13 | 18.5 | 76 | 0.05 | 0.81 | 0.18 | Immature |
| 900119000341576 | Crispant | Male | 1.59 | 39.06 | 59.35 | 38025 | 2018-02-13 | 13 | 27 | 0.26 | 1.29 | | |
| 900119000341282 | Control | Male | | | | | 2018-03-23 | 22.9 | 117 | 0.02 | 0.14 | 0.01 | Immature |
| 900119000341806 | Control | Male | | | | | 2018-03-23 | 25 | 164 | 0.03 | 0.05 | 0.02 | Immature |
| 900119000341916 | Control | Male | | | | | 2018-03-23 | 22.2 | 109 | 0.03 | 0.05 | 0.04 | Immature |
| 900119000341853 | Control | Male | | | | | 2018-03-23 | 29.3 | 321 | 1.26 | 2.98 | 9.69 | Mature |
| 900119000341547 | Control | Male | | | | | 2018-03-23 | 30 | 371 | 1.41 | 4.97 | 13.2 | Mature |
| 900119000341838 | Control | Male | | | | | 2018-03-23 | 28.9 | 305 | 1.54 | 2.92 | 3.74 | Mature |
| 900119000341664 | Control | Male | | | | | 2018-03-23 | 26 | 227 | 1.68 | 3.34 | 10.28 | Mature |
| 900119000341492 | Control | Male | | | | | 2018-03-23 | 25.6 | 231 | 1.8 | 4.34 | 22.34 | Mature |
| 900119000341490 | Control | Male | | | | | 2018-03-23 | 27.6 | 295 | 1.89 | 4.41 | 10.82 | Mature |
| 900119000341606 | Control | Male | | | | | 2018-03-23 | 23.2 | 162 | 2.06 | 2.65 | 20.82 | Mature |
| 900119000341121 | Control | Male | | | | | 2018-03-23 | 29 | 306 | 2.25 | 3.18 | | Mature |
| 900119000341997 | Control | Male | | | | | 2018-03-23 | 23.7 | 169 | 2.41 | 4.16 | 14.76 | Mature |
| 900119000341832 | Control | Male | | | | | 2018-03-23 | 26.3 | 239 | 2.49 | 3.55 | 19.53 | Mature |
| 900119000341443 | Control | Male | | | | | 2018-03-23 | 26.7 | 264 | 2.78 | 3.21 | 15.3 | Mature |
| 900119000341834 | Control | Male | | | | | 2018-03-23 | 27.9 | 297 | 2.79 | 4.21 | 17.48 | Mature |
| 900119000341772 | Control | Male | | | | | 2018-03-23 | 26.5 | 259 | 2.84 | 5.86 | 17.24 | Mature |
| 900119000341829 | Control | Male | | | | | 2018-03-23 | 27 | 249 | 3.36 | 3.63 | 22.71 | Mature |
| 900119000341767 | Crispant | Male | 4.12 | 13.83 | 82.05 | 39330 | 2018-03-23 | 23.5 | 143 | 0.05 | 0.61 | 0.16 | Immature |
| 900119000341889 | Crispant | Male | 0.29 | 19.77 | 79.93 | 41290 | 2018-03-23 | 30 | 342 | 1.84 | 4.92 | 7.46 | Mature |
| 900119000341699 | Crispant | Male | 19.15 | 18.74 | 62.1 | 39614 | 2018-03-23 | 27.5 | 255 | 1.89 | 1.92 | 12.46 | Mature |
| 900119000341273 | Crispant | Male | 14.41 | 22.12 | 63.46 | 36779 | 2018-03-23 | 15.7 | 54 | 7.17 | 1.17 | 12.79 | Mature |
| 900119000341452 | Crispant | Male | 20.58 | 24.61 | 54.81 | 37246 | 2018-03-23 | 20 | 83 | 0.04 | 0.05 | 0.03 | Immature |
| 900119000341578 | Crispant | Male | 8.01 | 16.62 | 75.38 | 39590 | 2018-03-23 | 24.7 | 168 | 0.04 | 0.12 | 0.06 | Immature |
| 900119000341839 | Crispant | Male | 23.9 | 18.54 | 57.57 | 30206 | 2018-03-23 | 22 | 112 | 0.04 | 0.05 | 0.04 | Immature |
| 900119000341921 | Crispant | Male | 13.86 | 13.93 | 72.21 | 36216 | 2018-03-23 | 22.2 | 118 | 0.04 | 0.28 | | Immature |
| 900119000341169 | Crispant | Male | 24.33 | 42.95 | 32.72 | 38378 | 2018-03-23 | 25.5 | 179 | 0.04 | 0.17 | 0.1 | Immature |
| 900119000341635 | Crispant | Male | 32.85 | 14.77 | 52.37 | 43885 | 2018-03-23 | 23.1 | 120 | 0.04 | 0.15 | 0.04 | Immature |
| 900119000341897 | Crispant | Male | 16.68 | 40.93 | 42.39 | 35835 | 2018-03-23 | 26.6 | 250 | 2.06 | 3.65 | 10.04 | Mature |
| 900119000341780 | Crispant | Male | 60.21 | 12.26 | 27.53 | 40156 | 2018-03-23 | 27.7 | 275 | 2.08 | 2.54 | 12.15 | Mature |
| 900119000341583 | Crispant | Male | 15.97 | 46.83 | 37.2 | 42196 | 2018-03-23 | 27.8 | 296 | 2.1 | 4.66 | 14.62 | Mature |
| 900119000341167 | Crispant | Male | 41.5 | 6.26 | 52.24 | 33433 | 2018-03-23 | 25 | 210 | 2.59 | 3.45 | 15.76 | Mature |
| 900119000341367 | Crispant | Male | 22.13 | 30.48 | 47.39 | 44515 | 2018-03-23 | 23.5 | 172 | 3.74 | 3.73 | 12.64 | Mature |
| 900119000341186 | Control | Male | | | | | 2018-09-25 | 33.2 | 489 | 5.19 | 18.08 | | Mature |
| 900119000341069 | Control | Male | | | | | 2018-09-25 | 35.5 | 555 | 7.59 | 17.84 | | Mature |
| 900119000341160 | Control | Male | | | | | 2018-09-25 | 38.5 | 751 | 7.16 | 12.6 | | Mature |
| 900119000341361 | Control | Male | | | | | 2018-09-25 | 32.4 | 461 | 9.09 | 22.04 | | Mature |
| 900119000343398 | Control | Male | | | | | 2018-09-25 | 36 | 564 | 5.09 | 18.75 | | Mature |
| 900119000341797 | Control | Male | | | | | 2018-09-25 | 40 | 723 | 0.05 | 0.46 | | Immature |
| 900119000341278 | Control | Male | | | | | 2018-09-25 | 38.7 | 683 | 8.58 | 13.16 | | Mature |
| 900119000341057 | Control | Male | | | | | 2018-09-25 | 32.2 | 405 | 5.91 | 39.4 | | Mature |
| 900119000341880 | Control | Male | | | | | 2018-09-25 | 34 | 458 | 7.36 | 16.96 | | Mature |
| 900119000341199 | Control | Male | | | | | 2018-09-25 | 34 | 491 | 5.68 | 16.38 | | Mature |
| 900119000341733 | Control | Male | | | | | 2018-09-25 | 32 | 440 | 8.5 | 29.96 | | Mature |
| 900119000341434 | Control | Male | | | | | 2018-09-25 | 29 | 330 | 6.06 | 36 | | Mature |
| 900119000341856 | Control | Male | | | | | 2018-09-25 | 31.2 | 416 | 6.85 | 25.7 | | Mature |
| 900119000341016 | Control | Male | | | | | 2018-09-25 | 28 | 289 | 6.8 | 14.06 | | Mature |
| 900119000341463 | Crispant | Male | 10.63 | 29.81 | 59.57 | 38621 | 2018-09-25 | 38.5 | 722 | 0.06 | 0.73 | | Immature |
| 900119000341953 | Crispant | Male | 36.3 | 34.83 | 28.87 | 37969 | 2018-09-25 | 44 | 995 | 0.06 | 0.65 | | Immature |
| 900119000341926 | Crispant | Male | 34.35 | 14.52 | 51.12 | 33181 | 2018-09-25 | 44.5 | 1030 | 0.08 | 0.97 | | Immature |
| 900119000341100 | Crispant | Male | 42.16 | 20.03 | 37.81 | 35119 | 2018-09-25 | 40.5 | 818 | 0.06 | 0.61 | | Immature |
| 900119000341099 | Crispant | Male | 35.62 | 27.36 | 37.02 | 44999 | 2018-09-25 | 33 | 501 | 9.57 | 12.04 | | Mature |
| 900119000341387 | Crispant | Male | 5.95 | 32.88 | 61.18 | 34790 | 2018-09-25 | 37.5 | 595 | 0.08 | 0.95 | | Immature |
| 900119000341602 | Crispant | Male | 7.51 | 20.52 | 71.97 | 34047 | 2018-09-25 | 43 | 978 | 0.05 | 0.87 | | Immature |

| | | | | | | | | | | | | |
|---|---|---|---|---|---|---|---|---|---|---|---|---|
| 900119000341978 | Crispant | Male | 59.05 | 4.52 | 36.43 | 32262 | 2018-09-25 | 39 | 757 | 0.09 | 0.99 | Immature |
| 900119000341949 | Crispant | Male | 27.88 | 29.51 | 42.61 | 41278 | 2018-09-25 | 22.2 | 163 | 11.82 | 11.81 | Mature |
| 900119000341336 | Control | Female | | | | | 2020-12-11 | 88 | 9870 | | | Mature |
| 900119000341209 | Control | Female | | | | | 2020-12-11 | 84 | 8775 | | | Mature |
| 900119000341459 | Control | Female | | | | | 2020-12-11 | 89 | 10875 | | | Mature |
| 900119000341927 | Control | Female | | | | | 2020-12-11 | 90 | 11340 | | | Mature |
| 900119000341152 | Control | Female | | | | | 2020-12-11 | 86 | 9245 | | | Mature |
| 900119000341197 | Control | Female | | | | | 2020-12-11 | 93 | 11390 | | | Mature |
| 900119000341869 | Control | Female | | | | | 2020-12-11 | 89 | 9485 | | | Mature |
| 900119000341518 | Control | Female | | | | | 2020-12-11 | 89 | 10815 | | | Mature |
| 900119000341268 | Control | Female | | | | | 2020-12-11 | 88 | 10755 | | | Mature |
| 900119000341248 | Control | Female | | | | | 2020-12-11 | 86 | 9755 | | | Mature |
| 900119000341211 | Control | Female | | | | | 2020-12-11 | 91 | 10250 | | | Mature |
| 900119000341454 | Control | Female | | | | | 2020-12-11 | 83 | 7010 | | | Mature |
| 900119000341076 | Control | Female | | | | | 2020-12-11 | 89 | 10870 | | | Mature |
| 900119000341711 | Control | Female | | | | | 2020-12-11 | 90 | 11505 | | | Mature |
| 900119000341909 | Control | Female | | | | | 2020-12-11 | 84 | 8440 | | | Mature |
| 900119000341342 | Control | Female | | | | | 2020-12-11 | 83 | 8555 | | | Mature |
| 900119000341421 | Control | Female | | | | | 2020-12-11 | 91 | 12460 | | | Mature |
| 900119000341073 | Control | Female | | | | | 2020-12-11 | 80 | 7105 | | | Mature |
| 900119000341088 | Control | Female | | | | | 2020-12-11 | 63 | 3318 | 0.30 | | Immature |
| 900119000341402 | Crispant | Female | 21.21 | 23.92 | 54.86 | 42831 | 2020-12-11 | 76 | 5344 | 0.45 | | Immature |
| 900119000341680 | Crispant | Female | 23.11 | 4.88 | 72.01 | 40106 | 2020-12-11 | 87 | 8770 | 0.78 | | Immature |
| 900119000341292 | Crispant | Female | 20.63 | 18.97 | 60.4 | 33008 | 2020-12-11 | 85 | 6465 | 0.71 | | Mature |
| 900119000341567 | Crispant | Female | 13.49 | 32.46 | 54.05 | 39580 | 2020-12-11 | 82 | 8045 | | | Mature |
| 900119000341605 | Crispant | Female | 31.37 | 16.08 | 52.55 | 49023 | 2020-12-11 | 85 | 9100 | | | Mature |
| 900119000341114 | Crispant | Female | 11.91 | 20.08 | 68.01 | 38186 | 2020-12-11 | 89 | 9955 | | | Mature |
| 900119000341138 | Crispant | Female | 65.08 | 7.36 | 27.56 | 39263 | 2020-12-11 | 93 | 10665 | | | Mature |
| 900119000341955 | Crispant | Female | 16.79 | 20.32 | 62.89 | 33525 | 2020-12-11 | 90 | 11490 | | | Mature |
| 900119000341370 | Crispant | Female | 31.43 | 13.19 | 55.39 | 42179 | 2020-12-11 | 87 | 10890 | | | Mature |
| 900119000341977 | Crispant | Female | 33.42 | 6.53 | 60.05 | 33963 | 2020-12-11 | 89 | 9290 | | | Mature |
| 900119000341597 | Crispant | Female | 42.99 | 8.35 | 48.67 | 39106 | 2020-12-11 | 91 | 12335 | | | Mature |
| 900119000341917 | Crispant | Female | 19.89 | 15.9 | 64.21 | 33585 | 2020-12-11 | 86 | 7940 | | | Mature |
| 900119000341014 | Crispant | Female | 37.56 | 12.01 | 50.43 | 39676 | 2020-12-11 | 87 | 9300 | | | Mature |
| 900119000341023 | Crispant | Female | 24.17 | 25.84 | 49.99 | 41805 | 2020-12-11 | 58 | 2420 | 0.49 | | Immature |
| 900119000341456 | Crispant | Female | | | | | 2020-12-11 | 95 | 11735 | 0.72 | | Immature |
| 900119000341645 | Crispant | Female | 38.68 | 22.22 | 39.09 | 28656 | 2020-12-11 | 85 | 8060 | | | Mature |
| 900119000341347 | Crispant | Female | 26.78 | 12.54 | 60.68 | 41865 | 2020-12-11 | 80 | 6775 | | | Mature |
| 900119000341877 | Crispant | Female | 0.11 | 21.38 | 78.52 | 34867 | 2020-12-11 | 78 | 6060 | 0.63 | | Immature |
| 900119000341864 | Crispant | Female | 11.49 | 9.03 | 79.48 | 30687 | 2020-12-11 | 82 | 7435 | | | Mature |
| 900119000341362 | Crispant | Female | 15.45 | 28.97 | 55.58 | 29755 | 2020-12-11 | 70 | 4584 | 0.41 | | Immature |

**Biometrics and genotypes for 2020 year-class**

| | | | 2022-06-17 | 2022-11-02 | 2023-09-27 | 2022-06-17 | | 2022-11-03 | | 2023-06-14 | |
|---|---|---|---|---|---|---|---|---|---|---|---|
| **PIT tag** | **Sex** | **Indel geno-type** | **Maturation status** | **Maturation status** | **Maturation status** | **Length (cm)** | **Weight (g)** | **Length (cm)** | **Weight (g)** | **Length (cm)** | **Weight (g)** |
| 900119000089429 | Male | FS/FS | Immature | Immature | | 28.5 | 335 | 375 | 876 | | |
| 900119000089161 | Male | FS/FS | Immature | Immature | | 24 | 167 | 345 | 596 | | |
| 04193FBCED | Male | FS/FS | Immature | Immature | | 31.3 | 391 | 465 | 1357 | | |
| 04179E0A5B | Male | FS/FS | Immature | Immature | | 24 | 211 | 350 | 817 | | |
| 900119000470075 | Male | FS/FS | Immature | Mature | | 31.5 | 417 | 455 | 1320 | | |
| 989001003401804 | Male | FS/FS | Immature | | | 25.5 | 188 | | | | |
| 900119000089567 | Male | FS/FS | Immature | | | 27.5 | 240 | | | | |
| 900119000089352 | Male | FS/FS | Immature | | | 29.5 | 343 | | | | |
| 989001003401981 | Male | FS/WT | Immature | Immature | | 27.5 | 265 | 410 | 1009 | | |
| 900212000069405 | Male | FS/WT | Immature | Immature | | 29 | 300 | 425 | 1085 | | |
| 900119000089711 | Male | FS/WT | Immature | Immature | | 28.5 | 282 | 420 | 1009 | | |
| 900119000470488 | Male | FS/WT | Mature | Mature | | 30.5 | 351 | 420 | 1002 | | |
| 041A5A836D | Male | FS/WT | Immature | Mature | | 31.2 | 412 | 430 | 1018 | | |
| 900119000470941 | Male | FS/WT | Mature | Mature | | 29 | 297 | 340 | 504 | | |
| 0417212E69 | Male | FS/WT | Mature | Mature | | 28 | 261 | 340 | 492 | | |
| 900119000042555 | Male | FS/WT | Mature | Mature | | 32 | 385 | 400 | 861 | | |
| 0419E5E473 | Male | FS/WT | Mature | Mature | | 34.5 | 464 | 410 | 908 | | |
| 900119000342749 | Male | FS/WT | Mature | Mature | | 28.5 | 280 | 340 | 544 | | |
| 0419E64AF1 | Male | FS/WT | Immature | Mature | | 29.3 | 318 | 410 | 933 | | |
| 041A5A82A3 | Male | FS/WT | Mature | Mature | | 26.5 | 251 | 325 | 492 | | |
| 0419E64F35 | Male | FS/WT | Mature | Mature | | 17.2 | 67 | 210 | 123 | | |
| 989001003400840 | Male | FS/WT | Mature | Mature | | 15.5 | 48 | 190 | 100 | | |
| 0419E6335A | Male | FS/WT | | Mature | | | | 340 | 469 | | |
| 041A5B727B | Male | FS/WT | Immature | Mature | | 27.5 | 292 | 350 | 670 | | |
| 900119000088681 | Male | FS/WT | Mature | Mature | | 32 | 372 | 355 | 560 | | |
| 900119000066295 | Male | FS/WT | Mature | Mature | | 41 | 857 | 455 | 1126 | | |
| 900119000344037 | Male | FS/WT | Mature | Mature | | 31 | 399 | 355 | 578 | | |
| 985170001756571 | Male | FS/WT | Mature | Mature | | 31.5 | 410 | 375 | 705 | | |
| 900119000066162 | Male | FS/WT | Mature | Mature | | 33 | 484 | 390 | 778 | | |
| 989001003399391 | Male | FS/WT | Immature | | | 31.5 | 380 | | | | |
| 900119000088969 | Male | FS/WT | Immature | | | 32 | 396 | | | | |
| 900119000089358 | Male | WT/WT | Immature | Immature | | 29.2 | 322 | 400 | 884 | | |
| 985170001951304 | Male | WT/WT | Mature | Mature | | 29.5 | 371 | 385 | 950 | | |
| 900119000089275 | Male | WT/WT | Immature | Mature | | 33.3 | 509 | 450 | 1309 | | |
| 0419E65959 | Male | WT/WT | Immature | Mature | | 29 | 326 | 420 | 1122 | | |
| 900119000066682 | Male | WT/WT | Mature | Mature | | 35.5 | 506 | 425 | 983 | | |
| 900119000043032 | Male | WT/WT | Mature | Mature | | 32 | 410 | 380 | 707 | | |
| 985170001991176 | Male | WT/WT | Mature | Mature | | 32 | 391 | 390 | 747 | | |
| 900119000042236 | Male | WT/WT | | Mature | | 35 | 540 | 400 | 764 | | |
| 992002000236708 | Male | WT/WT | Mature | Mature | | 30.5 | 458 | 360 | 612 | | |
| 989001003400014 | Male | WT/WT | Mature | Mature | | 31.5 | 441 | 360 | 616 | | |
| 900119000042916 | Male | WT/WT | Immature | | | 36 | 536 | | | | |
| 0006E8F2AD | Male | WT/WT | Immature | | | 22 | 186 | | | | |
| 985170001867223 | Female | FS/FS | | | Immature | | | | | 49 | 1450 |
| 900119000066042 | Female | FS/FS | | | Immature | | | | | 71.5 | 5052 |
| 900119000047689 | Female | FS/FS | | | Immature | | | | | 56.8 | 2100 |
| 041A09D411 | Female | FS/FS | | | Immature | | | | | 58.5 | 2670 |
| 992002000252479 | Female | FS/WT | | | Immature | | | | | 66.5 | 3744 |
| 985170001944435 | Female | FS/WT | | | Immature | | | | | 61 | 3146 |
| 985170001907443 | Female | FS/WT | | | Immature | | | | | 63 | 2976 |
| 985170001820944 | Female | FS/WT | | | Immature | | | | | 65.4 | 4268 |
| 900212000200094 | Female | FS/WT | | | Immature | | | | | 63.2 | 3498 |
| 900119000470156 | Female | FS/WT | | | Immature | | | | | 64 | 3498 |
| 900119000341861 | Female | FS/WT | | | Mature | | | | | 61.9 | 3560 |
| 900119000341584 | Female | FS/WT | | | Mature | | | | | 66.8 | 4302 |
| 900119000089795 | Female | FS/WT | | | Mature | | | | | 67.7 | 4680 |
| 900119000089593 | Female | FS/WT | | | Mature | | | | | 65.7 | 4056 |
| 900119000088947 | Female | FS/WT | | | Immature | | | | | 57 | 2350 |
| 900119000088023 | Female | FS/WT | | | Immature | | | | | 66.5 | 3776 |
| 900119000066933 | Female | FS/WT | | | Mature | | | | | 55.9 | 2698 |
| 900119000066744 | Female | FS/WT | | | Mature | | | | | 64.6 | 3972 |
| 900119000066552 | Female | FS/WT | | | Immature | | | | | 48 | 1406 |
| 900119000042694 | Female | FS/WT | | | Immature | | | | | 63.5 | 3628 |
| 900119000040357 | Female | FS/WT | | | Mature | | | | | 60.4 | 3230 |
| 041A5B7298 | Female | FS/WT | | | Immature | | | | | 62.2 | 2840 |

| | | | | | | | | | |
|---|---|---|---|---|---|---|---|---|---|
| 041A5B6CE0 | Female | FS/WT | | | Mature | | | 71.2 | 5374 |
| 041A311460 | Female | FS/WT | | | Immature | | | 55 | 2610 |
| 041A0439AA | Female | FS/WT | | | Mature | | | 66.4 | 4164 |
| 0419E66A2C | Female | FS/WT | | | Mature | | | 69.1 | 5280 |
| 0006E8E0E4 | Female | FS/WT | | | Mature | | | 69 | 4992 |
| 992002000257730 | Female | FS/WT | | | Mature | | | 64.5 | 4160 |
| 900119000089733 | Female | FS/WT | | | Mature | | | 66.7 | 4332 |
| 900119000066463 | Female | FS/WT | | | Mature | | | 61.9 | 4090 |
| 900119000066440 | Female | FS/WT | | | Immature | | | 71.5 | 4292 |
| 041A09CE82 | Female | FS/WT | | | Immature | | | 54 | 2876 |
| 0419E669D0 | Female | FS/WT | | | Mature | | | 65.2 | 4398 |
| 0419E6161E | Female | FS/WT | | | Immature | | | 64 | 3006 |
| 0419E614DC | Female | FS/WT | | | Mature | | | 68.7 | 4848 |
| 989001003920292 | Female | WT/WT | | | Mature | | | 67.3 | 4710 |
| 900119000089385 | Female | WT/WT | | | Mature | | | 63.2 | 4130 |
| 900119000042644 | Female | WT/WT | | | Immature | | | 62.2 | 3014 |
| 989001003399569 | Female | WT/WT | | | Mature | | | 69.5 | 4570 |
| 04185C2DEE | Female | WT/WT | | | Mature | | | 63.9 | 3800 |
| 000630ED7A | Female | WT/WT | | | Mature | | | 70.5 | 5072 |
| 989001003400588 | Female | WT/WT | | | Mature | | | 69.5 | 4958 |
| 982000407900306 | Female | WT/WT | | | Mature | | | 74.3 | 6250 |
| 900119000089441 | Female | WT/WT | | | Mature | | | 68.3 | 4748 |

Biometrics and genotypes for 2021 year-class

| PIT tag | Sex | Natural geno-type | Indel geno-type | Combined genotype | 2023-03-22 | 2023-04-24 | 2023-11-03 | 2024-10-28, 2024-11-07 | 2023-03-22 | | 2023-04-24 | | 2023-11-03 | | 2024-10-28, 2024-11-07 | |
|---|---|---|---|---|---|---|---|---|---|---|---|---|---|---|---|---|
| | | | | | 11-KT (ng/mL) | Maturation status | Maturation status | Maturation status | Length (cm) | Weight (g) | Length (cm) | Weight (g) | Length (cm) | Weight (g) | Length (cm) | Weight (g) |
| 041A5B6A6C | Male | EL | FS/FS | FS/FS | 0.01 | Immature | Immature | | 41 | 776 | 44 | 1080 | 68 | 4782 | | |
| 900119000066642 | Male | EL | FS/FS | FS/FS | 2.44 | Immature | Immature | | 31.2 | 372 | 35.1 | 575 | 54 | 2411 | | |
| 041A099614 | Male | EL | FS/FS | FS/FS | 0.64 | Immature | Immature | | 38.9 | 832 | 42.5 | 1115 | 65 | 4315 | | |
| 0419E66904 | Male | EL | FS/FS | FS/FS | 1.05 | Immature | Immature | | 31 | 386 | 35.1 | 605 | 56 | 2490 | | |
| 900119000089865 | Male | EL | FS/FS | FS/FS | 1.28 | Immature | Immature | | 44 | 1148 | 49.3 | 1665 | 61 | 2040 | | |
| 989001003400610 | Male | EL | FS/FS | FS/FS | 5.98 | Immature | Immature | | 38.2 | 774 | 41.5 | 1020 | 64 | 4310 | | |
| 900119000066368 | Male | EE | FS/FS | FS/FS | 1.19 | Immature | Immature | | 40.8 | 996 | 43.9 | 1260 | 63 | 4252 | | |
| 900119000088221 | Male | EL | FS/FS | FS/FS | 2.08 | Immature | Immature | | 40.6 | 910 | 44.4 | 1185 | 64 | 4462 | | |
| 900119000471020 | Male | EE | FS/FS | FS/FS | 0.65 | Immature | Immature | | 40.4 | 840 | 44.1 | 1140 | 65 | 3783 | | |
| 989001003400880 | Male | EL | FS/FS | FS/FS | 1.36 | Immature | Immature | | 41.6 | 1024 | 45.4 | 1335 | 66 | 4600 | | |
| 900119000358614 | Male | EL | FS/FS | FS/FS | 1.07 | Immature | Immature | | 41 | 852 | 44.3 | 1135 | 61 | 3130 | | |
| 900119000414780 | Male | EE | FS/FS | FS/FS | 5.47 | Immature | Immature | | 47 | 1374 | 52 | 1840 | 74 | 5688 | | |
| 900119000343958 | Male | EL | FS/FS | FS/FS | 3.5 | Immature | Immature | | 42 | 994 | 46.4 | 1400 | 68 | 4747 | | |
| 041A725FD1 | Male | EL | FS/FS | FS/FS | 1.35 | Immature | Immature | | 41.2 | 884 | 45.1 | 1195 | 64 | 3577 | | |
| 900212000069302 | Male | EE | FS/FS | FS/FS | 2.69 | Immature | Immature | | 44 | 1200 | 48 | 1570 | 67 | 4790 | | |
| 900119000047205 | Male | EL | FS/FS | FS/FS | 5.79 | Immature | Immature | | 37.1 | 678 | 41.9 | 1000 | 72 | 3450 | | |
| 900119000342344 | Male | | FS/FS | FS/FS | 1.57 | Immature | Immature | | 35.2 | 588 | 38.5 | 800 | 56 | 2670 | | |
| 900119000089694 | Male | EL | FS/FS | FS/FS | 2.17 | Immature | Immature | | 33.3 | 500 | 36.3 | 665 | 52 | 2280 | | |
| 900119000042141 | Male | EL | FS/FS | FS/FS | 1.73 | Immature | Immature | | 39.6 | 842 | 43.5 | 1140 | 60 | 3206 | | |
| 041A711A52 | Male | EL | FS/FS | FS/FS | 3.13 | Immature | Immature | | 40.6 | 850 | 44.2 | 1110 | 61 | 3000 | | |
| 04193F8586 | Male | EL | FS/FS | FS/FS | 3.25 | Immature | Immature | | 41.5 | 946 | 45.6 | 1300 | 66 | 4505 | | |
| 041A72715B | Male | EL | FS/FS | FS/FS | 1.32 | Immature | Immature | | 38.8 | 762 | 43 | 1110 | 61 | 3170 | | |
| 900119000042473 | Male | EL | FS/FS | FS/FS | 2.32 | Immature | Immature | | 36.7 | 626 | 40.8 | 900 | 57 | 2666 | | |
| 900119000047636 | Male | EL | FS/FS | FS/FS | 0.01 | Immature | Immature | | 39.5 | 754 | 44.1 | 1135 | 68 | 4830 | | |
| 900119000066739 | Male | LL | FS/FS | FS/FS | 1.22 | Immature | Immature | | 33.3 | 466 | 36.5 | 635 | 56 | 2430 | | |
| 041808028E | Male | EL | FS/FS | FS/FS | 1.94 | Immature | Immature | | 37.7 | 662 | 42 | 920 | 59 | 2549 | | |
| 900119000341566 | Male | EL | FS/FS | FS/FS | 1.26 | Mature | Immature | | 40.8 | 896 | 45.4 | 1320 | 68 | 4936 | | |
| 900119000088150 | Male | EL | FS/FS | FS/FS | 3.92 | Immature | Immature | | 35.6 | 612 | 39 | 790 | 63 | 3870 | | |
| 900119000089686 | Male | EL | FS/FS | FS/FS | 1.38 | Immature | Immature | | 37.2 | 767 | 38.5 | 635 | 61 | 3564 | | |
| 041818A39D | Male | EL | FS/FS | FS/FS | 0.94 | Immature | Immature | | 30.2 | 372 | 33 | 520 | 49 | 1782 | | |
| 900212000184618 | Male | EL | FS/FS | FS/FS | 1.45 | Immature | Immature | | 37 | 704 | 40.8 | 950 | 55 | 2444 | | |
| 900119000089925 | Male | EL | FS/FS | FS/FS | 3.83 | Immature | Immature | | 37 | 636 | 39.1 | 795 | 60 | 3170 | | |
| 989001003399570 | Male | EE | FS/FS | FS/FS | 3.39 | Immature | Immature | | 43.3 | 1034 | 47.2 | 1340 | 57 | 2722 | | |
| 041A7117CA | Male | EL | FS/FS | FS/FS | 14.51 | Mature | Mature | | 38.4 | 738 | 42 | 990 | 56 | 2650 | | |
| 985170001974368 | Male | EL | FS/FS | FS/FS | 10.88 | Mature | Mature | | 38.2 | 688 | 42 | 965 | 54 | 2095 | | |
| 900119000342423 | Male | EE | FS/FS | FS/FS | 5.62 | Mature | Mature | | 44 | 1094 | 48.4 | 1375 | 64 | 3440 | | |
| 0418C7A798 | Male | EL | FS/FS | FS/FS | 3.36 | Immature | Mature | | 34 | 512 | 36.8 | 645 | 54 | 1984 | | |
| 041A7276C6 | Male | EL | FS/FS | FS/FS | 8.18 | Mature | Mature | | 40 | 812 | 44 | 1070 | 64 | 3771 | | |
| 900119000089455 | Male | EL | FS/FS | FS/FS | 1.69 | Immature | Mature | | 36.2 | 528 | 37.1 | 590 | 54 | 2319 | | |
| 985170001907354 | Male | EE | FS/FS | FS/FS | 19.32 | Mature | Mature | | 37.7 | 670 | 40.2 | 820 | 45 | 1219 | | |
| 900119000066703 | Male | EL | FS/FS | FS/FS | 4.11 | Immature | Mature | | 37.6 | 722 | 42.2 | 1030 | 49 | 3035 | | |
| 041A7193FB | Male | EL | FS/FS | FS/FS | 3.89 | Mature | Mature | | 33.6 | 524 | 37 | 755 | 46 | 1611 | | |
| 985170001807861 | Male | EL | FS/FS | FS/FS | 12.37 | Mature | Mature | | 36.4 | 588 | 39.1 | 840 | 44 | 993 | | |
| 041A719426 | Male | EL | FS/FS | FS/FS | 10.56 | Mature | Mature | | 41.4 | 946 | 46.3 | 1430 | 66 | 5067 | | |
| 900119000066021 | Male | LL | FS/FS | FS/FS | 23.4 | Mature | Mature | | 38.8 | 752 | 41 | 950 | 44 | 1172 | | |
| 900119000042931 | Male | EE | FS/FS | FS/FS | 8.34 | Mature | Mature | | 36.4 | 668 | 38.6 | 810 | 48 | 1578 | | |
| 04193FBDA4 | Male | EL | FS/FS | FS/FS | 3.35 | Immature | | | 42 | 930 | 46 | 1235 | | | | |
| 900119000414801 | Male | LL | FS/FS | FS/FS | 0.66 | Immature | | | 29.5 | 340 | 32 | 420 | | | | |
| 900119000089114 | Male | LL | FS/FS | FS/FS | 1.72 | Immature | | | 36 | 600 | 39.3 | 760 | | | | |
| 041818840B | Male | EL | FS/FS | FS/FS | 2.1 | Immature | | | 41 | 826 | 44.6 | 1155 | | | | |
| 900119000047375 | Male | EL | FS/FS | FS/FS | 0.95 | Immature | | | 31.5 | 394 | 33.8 | 500 | | | | |
| 985170001856947 | Male | EL | FS/FS | FS/FS | 0.81 | Immature | | | 39 | 794 | 43 | 1105 | | | | |
| 0419E61548 | Male | EL | FS/FS | FS/FS | 6.12 | Immature | | | 34.8 | 590 | 38.5 | 810 | | | | |
| 985170001948511 | Male | EL | FS/FS | FS/FS | 1.99 | Immature | | | 43 | 1092 | 47.5 | 1510 | | | | |
| 900119000088993 | Male | EE | FS/FS | FS/FS | 1.03 | Immature | | | 45.5 | 1316 | 50.2 | 1965 | | | | |
| 04193FA3AD | Male | | FS/FS | FS/FS | 2.92 | Immature | | | 41.5 | 890 | 46 | 1265 | | | | |
| 989001003401483 | Male | EE | FS/FS | FS/FS | 2.31 | Immature | | | 38 | 682 | 40.5 | 780 | | | | |
| 989001003401227 | Male | EL | FS/FS | FS/FS | 1.85 | Immature | | | 35.6 | 540 | 38 | 710 | | | | |
| 900119000066835 | Male | EL | FS/FS | FS/FS | 0.68 | Immature | | | 44.8 | 1376 | 49 | 1875 | | | | |
| 0419E61D84 | Male | EL | FS/FS | FS/FS | 14.27 | Mature | | | 36 | 604 | 39.4 | 860 | | | | |
| 041A099FEA | Male | EL | FS/FS | FS/FS | 6.61 | Mature | | | 38.1 | 686 | 41 | 845 | | | | |
| 041A711A56 | Male | EE | FS/FS | FS/FS | 6.64 | Mature | | | 40.6 | 806 | 43.4 | 1030 | | | | |
| 041A045652 | Male | EL | FS/WT | FS/E | 15.82 | Mature | Mature | | 41 | 866 | 44 | 1110 | 48 | 1324 | | |
| 989001003401770 | Male | EL | FS/WT | FS/E | 11.38 | Mature | Mature | | 45 | 1336 | 50 | 1900 | 59 | 2973 | | |
| 0416F1E3A7 | Male | EL | FS/WT | FS/E | 13.04 | Mature | Mature | | 37.2 | 708 | 40.5 | 915 | 51 | 2090 | | |
| 900119000047905 | Male | EE | FS/WT | FS/E | 3.95 | Mature | Mature | | 48.5 | 1546 | 53 | 2060 | 65 | 3676 | | |
| 989001003400854 | Male | EL | FS/WT | FS/E | 12.01 | Mature | Mature | | 48.5 | 1628 | 52 | 1860 | 61 | 3606 | | |
| 041A045DFF | Male | EL | FS/WT | FS/E | 9.59 | Mature | Mature | | 45 | 1436 | 49.5 | 1970 | 56 | 2188 | | |
| 989001003401280 | Male | EE | FS/WT | FS/E | 12.28 | Mature | Mature | | 45.5 | 1176 | 49.8 | 1675 | 56 | 1946 | | |
| 0418C7C703 | Male | EE | FS/WT | FS/E | 22.3 | Mature | Mature | | 38 | 668 | 39.5 | 735 | 44 | 1205 | | |
| 041A5AA56C | Male | EL | FS/WT | FS/E | 17.19 | Mature | Mature | | 46.4 | 1372 | 50.5 | 1785 | 57 | 2315 | | |
| 992002000257134 | Male | EL | FS/WT | FS/E | 10.88 | Mature | Mature | | 45.2 | 1342 | 50 | 1880 | 63 | 4365 | | |
| 900119000088134 | Male | EL | FS/WT | FS/E | 6.49 | Mature | Mature | | 45 | 1228 | 50.1 | 1830 | 57 | 2160 | | |
| 900119000042543 | Male | EL | FS/WT | FS/E | 10.11 | Mature | Mature | | 41 | 964 | 44.5 | 1255 | 53 | 1657 | | |
| 041A043618 | Male | EL | FS/WT | FS/E | 20.87 | Mature | | | 42 | 1022 | 44.6 | 1145 | | | | |
| 900119000066036 | Male | EL | FS/WT | FS/E | 16.08 | Mature | | | 41 | 874 | 45.4 | 1150 | | | | |
| 041A3107D3 | Male | EL | FS/WT | FS/E | 22.61 | Mature | | | 44 | 1066 | 46.5 | 1325 | | | | |
| 992002000257908 | Male | EL | FS/WT | FS/E | 12.63 | Mature | | | 44 | 1186 | 47.5 | 1520 | | | | |
| 041A0454E5 | Male | EL | FS/WT | FS/E | 6.85 | Mature | | | 33.2 | 498 | 35.2 | 565 | | | | |
| 041917AECA | Male | EL | FS/WT | FS/E | 5.33 | Mature | | | 42 | 1042 | 45 | 1245 | | | | |
| 900119000042422 | Male | EL | FS/WT | FS/L | 6.19 | Immature | Immature | | 45.4 | 1316 | 50.1 | 1865 | 69 | 5075 | | |
| 900119000089310 | Male | EL | FS/WT | FS/L | 1.53 | Immature | Immature | | 40 | 848 | 43.2 | 1135 | 63 | 3702 | | |
| 900119000088458 | Male | LL | FS/WT | FS/L | 6.67 | Mature | Mature | | 32.6 | 454 | 36.2 | 635 | 46 | 1411 | | |
| 900119000343361 | Male | EL | FS/WT | FS/L | 36.32 | Mature | Mature | | 40.5 | 878 | 42.5 | 1010 | 45 | 1105 | | |
| 900119000042264 | Male | EL | FS/WT | FS/L | 12.52 | Mature | Mature | | 39.5 | 804 | 42.5 | 1020 | 47 | 1290 | | |
| 041A5AA4B2 | Male | LL | FS/WT | FS/L | 11.34 | Mature | Mature | | 38.7 | 764 | 42 | 1040 | 52 | 2175 | | |
| 900119000088343 | Male | EL | FS/WT | FS/L | 22.11 | Mature | Mature | | 41.5 | 970 | 45.5 | 1355 | 40 | 1582 | | |
| 989001003399371 | Male | LL | FS/WT | FS/L | 14.74 | Mature | Mature | | 39.5 | 858 | 43 | 1110 | 44 | 2340 | | |
| 900119000047247 | Male | LL | FS/WT | FS/L | 23.27 | Mature | Mature | | 42.8 | 1072 | 45 | 1170 | 51 | 1505 | | |
| 0419E65D30 | Male | LL | FS/WT | FS/L | 3.5 | Mature | Mature | | 30.6 | 388 | 34.5 | 515 | 43 | 1159 | | |
| 04185C502D | Male | LL | FS/WT | FS/L | 0.85 | Immature | Mature | | 31.1 | 418 | 34 | 560 | 43 | 1140 | | |
| 041A04437D | Male | LL | FS/WT | FS/L | 10.2 | Mature | Mature | | 38.2 | 738 | 42 | 1010 | 47 | 1080 | | |
| 000702E8D3 | Male | LL | FS/WT | FS/L | 10.93 | Mature | Mature | | 37.2 | 702 | 41.2 | 950 | 53 | 2082 | | |
| 900119000041482 | Male | EL | FS/WT | FS/L | 12.99 | Mature | Mature | | 36.6 | 652 | 40.5 | 935 | 47 | 1682 | | |
| 900119000344184 | Male | LL | FS/WT | FS/L | 10.65 | Mature | Mature | | 34.5 | 508 | 37 | 655 | 41 | 823 | | |
| 041A725CC1 | Male | LL | FS/WT | FS/L | 7.28 | Mature | Mature | | 45 | 1222 | 47.9 | 1535 | 57 | 2850 | | |
| 900119000343744 | Male | LL | FS/WT | FS/L | 7.01 | Mature | Mature | | 43 | 1156 | 46.8 | 1500 | 53 | 1930 | | |

| | | | | | | | | | | | | | | | | |
|---|---|---|---|---|---|---|---|---|---|---|---|---|---|---|---|---|
| 0419E662E5 | Male | EL | FS/WT | FS/L | 0.53 | Immature | | | 34.8 | 456 | 35.4 | 395 | | | | |
| 0418188561 | Male | EL | FS/WT | FS/L | 14.54 | Mature | | | 41.3 | 910 | 45.5 | 1230 | | | | |
| 900119000412840 | Male | LL | FS/WT | FS/L | 9.26 | Mature | | | 37.4 | 738 | 41.1 | 1015 | | | | |
| 0419E6419B | Male | LL | FS/WT | FS/L | 13.41 | Mature | | | 32.2 | 456 | 34 | 505 | | | | |
| 985170001955345 | Male | LL | FS/WT | FS/L | 14.2 | Mature | | | 43.8 | 1136 | 47.4 | 1585 | | | | |
| 000702DA43 | Male | LL | FS/WT | FS/L | 15.72 | Mature | | | 39.4 | 828 | 43.5 | 1165 | | | | |
| 989001003399826 | Male | LL | FS/WT | FS/L | 15.48 | Mature | | | 35.5 | 616 | 38.2 | 830 | | | | |
| 989001003400116 | Male | LL | FS/WT | FS/L | 19.74 | Mature | | | 43.2 | 1186 | 46.5 | 1505 | | | | |
| 041A310EE3 | Male | LL | FS/WT | FS/L | 9.54 | Mature | | | 40.3 | 845 | 43.5 | 1050 | | | | |
| 900119000042703 | Male | LL | FS/WT | FS/L | 9.67 | Mature | | | 45.2 | 1296 | 49.5 | 1665 | | | | |
| 900119000343476 | Male | LL | FS/WT | FS/L | 6.56 | Mature | | | 36.3 | 620 | 39.2 | 860 | | | | |
| 989001003400475 | Male | EL | WT/WT | EL | 22.66 | Mature | Mature | | 42.9 | 1028 | 45 | 1210 | 53 | 2157 | | |
| 900119000344223 | Male | EL | WT/WT | EL | 3.52 | Mature | Mature | | 36 | 586 | 38.5 | 705 | 45 | 1230 | | |
| 985170001716090 | Male | LL | WT/WT | LL | 0.01 | Immature | Immature | | 40.1 | 740 | 42.7 | 1030 | 62 | 3190 | | |
| 041A725C77 | Male | LL | WT/WT | LL | 6.52 | Mature | Mature | | 40.3 | 854 | 43.5 | 1085 | 51 | 1877 | | |
| 900119000342071 | Male | LL | WT/WT | LL | 10.03 | Mature | Mature | | 41 | 894 | 43.5 | 1155 | 48 | 1213 | | |
| 900119000066172 | Female | EL | FS/FS | FS/FS | | | | Immature | | | | | | | 79.5 | 6432 |
| 985170001924126 | Female | EL | FS/FS | FS/FS | | | | Immature | | | | | | | 78 | 5763 |
| 900119000089367 | Female | EE | FS/FS | FS/FS | | | | Immature | | | | | | | 79 | 6936 |
| 900119000066716 | Female | EL | FS/FS | FS/FS | | | | Immature | | | | | | | 78 | 5801 |
| 900119000088883 | Female | EL | FS/FS | FS/FS | | | | Immature | | | | | | | 76 | 5657 |
| 900119000342483 | Female | EL | FS/FS | FS/FS | | | | Immature | | | | | | | 85 | 8080 |
| 041A044002 | Female | EL | FS/FS | FS/FS | | | | Immature | | | | | | | 81 | 6770 |
| 041A711635 | Female | EL | FS/FS | FS/FS | | | | Immature | | | | | | | 86 | 9335 |
| 900119000089606 | Female | EL | FS/FS | FS/FS | | | | Immature | | | | | | | 80 | 6811 |
| 989001003163435 | Female | EL | FS/FS | FS/FS | | | | Immature | | | | | | | 88 | 10432 |
| 985170001836968 | Female | EL | FS/FS | FS/FS | | | | Immature | | | | | | | 81 | 7522 |
| 041A723072 | Female | EL | FS/FS | FS/FS | | | | Immature | | | | | | | 85 | 7199 |
| 900119000047167 | Female | EL | FS/FS | FS/FS | | | | Immature | | | | | | | 81.5 | 7250 |
| 0418188A33 | Female | EE | FS/FS | FS/FS | | | | Immature | | | | | | | 78 | 7034 |
| 985170001825551 | Female | EL | FS/FS | FS/FS | | | | Immature | | | | | | | 84 | 7472 |
| 900119000341676 | Female | EL | FS/FS | FS/FS | | | | Mature | | | | | | | 76 | 6662 |
| 989001003402200 | Female | EE | FS/FS | FS/FS | | | | Mature | | | | | | | 91 | 12996 |
| 04193F7D5D | Female | EL | FS/FS | FS/FS | | | | Mature | | | | | | | 79 | 6860 |
| 900119000088722 | Female | EL | FS/FS | FS/FS | | | | Mature | | | | | | | 79 | 7450 |
| 041A042643 | Female | EL | FS/FS | FS/FS | | | | Mature | | | | | | | 82 | 6930 |
| 900212000056889 | Female | LL | FS/FS | FS/FS | | | | Mature | | | | | | | 76 | 7096 |
| 041A719210 | Female | EL | FS/FS | FS/FS | | | | Mature | | | | | | | 73 | 7270 |
| 041A71140D | Female | EL | FS/FS | FS/FS | | | | Mature | | | | | | | 72 | 4686 |
| 900119000454376 | Female | EL | FS/FS | FS/FS | | | | Mature | | | | | | | 84 | 9784 |
| 04185C1EB2 | Female | EL | FS/FS | FS/FS | | | | Mature | | | | | | | 79 | 7980 |
| 04193FACDB | Female | EL | FS/FS | FS/FS | | | | Mature | | | | | | | 78 | 7270 |
| 900119000066901 | Female | EL | FS/FS | FS/FS | | | | Mature | | | | | | | 74 | 5494 |
| 989001003401800 | Female | EL | FS/FS | FS/FS | | | | Mature | | | | | | | 77 | 5870 |
| 041A726143 | Female | EL | FS/FS | FS/FS | | | | Mature | | | | | | | 77 | 6032 |
| 041A725905 | Female | EL | FS/FS | FS/FS | | | | Mature | | | | | | | 80 | 7562 |
| 900212000069371 | Female | EE | FS/FS | FS/FS | | | | Mature | | | | | | | 77 | 6584 |
| 041A09BE13 | Female | EL | FS/FS | FS/FS | | | | Mature | | | | | | | 66.5 | 3326 |
| 900119000047338 | Female | EL | FS/FS | FS/FS | | | | Mature | | | | | | | 82.5 | 8419 |
| 0419E65962 | Female | EL | FS/FS | FS/FS | | | | Mature | | | | | | | 96 | 14696 |
| 989001003401172 | Female | EL | FS/FS | FS/FS | | | | Mature | | | | | | | 80 | 6367 |
| 900119000342176 | Female | EL | FS/FS | FS/FS | | | | Mature | | | | | | | 79 | 8514 |
| 900119000342604 | Female | EL | FS/FS | FS/FS | | | | Mature | | | | | | | 73 | 5907 |
| 041A04324B | Female | EE | FS/FS | FS/FS | | | | Mature | | | | | | | 91 | 11619 |
| 900119000344563 | Female | EE | FS/FS | FS/FS | | | | Mature | | | | | | | 86.5 | 10720 |
| 900119000047863 | Female | EL | FS/FS | FS/FS | | | | Mature | | | | | | | 82 | 9780 |
| 0419407622 | Female | EE | FS/FS | FS/FS | | | | Mature | | | | | | | 83.5 | 9240 |
| 989001003399534 | Female | EE | FS/WT | FS/E | | | | Immature | | | | | | | 82 | 8989 |
| 041A7260FF | Female | EL | FS/WT | FS/L | | | | Immature | | | | | | | 81.5 | 8403 |
| 989001003399327 | Female | EL | FS/WT | FS/E | | | | Mature | | | | | | | 79 | 7634 |
| 985170001820427 | Female | EL | FS/WT | FS/E | | | | Mature | | | | | | | 85 | 10138 |
| 900119000088345 | Female | EL | FS/WT | FS/E | | | | Mature | | | | | | | | |
| 989001003401979 | Female | EL | FS/WT | FS/E | | | | Mature | | | | | | | 79 | 6582 |
| 0419E652CD | Female | EL | FS/WT | FS/E | | | | Mature | | | | | | | | |
| 041A719A4C | Female | EL | FS/WT | FS/E | | | | Mature | | | | | | | 87 | 9872 |
| 900119000470787 | Female | EL | FS/WT | FS/E | | | | Mature | | | | | | | 75 | 6470 |
| 0419E64FEE | Female | EL | FS/WT | FS/E | | | | Mature | | | | | | | 80 | 7058 |
| 04185C1C24 | Female | EL | FS/WT | FS/E | | | | Mature | | | | | | | 85 | 9242 |
| 900119000066787 | Female | EE | FS/WT | FS/E | | | | Mature | | | | | | | 77 | 6964 |
| 900119000066158 | Female | EL | FS/WT | FS/E | | | | Mature | | | | | | | | |
| 900119000412193 | Female | EL | FS/WT | FS/E | | | | Mature | | | | | | | | |
| 992002000259034 | Female | EL | FS/WT | FS/E | | | | Mature | | | | | | | | |
| 900119000066503 | Female | EE | FS/WT | FS/E | | | | Mature | | | | | | | 77 | 7120 |
| 900119000342854 | Female | EL | FS/WT | FS/E | | | | Mature | | | | | | | 84 | 9400 |
| 0419E5F1FA | Female | LL | FS/WT | FS/L | | | | Mature | | | | | | | 78 | 6482 |
| 900119000089641 | Female | LL | FS/WT | FS/L | | | | Mature | | | | | | | 80 | 7522 |
| 985170001749550 | Female | EL | FS/WT | FS/L | | | | Mature | | | | | | | 74 | 5068 |
| 900212000200484 | Female | EL | FS/WT | FS/L | | | | Mature | | | | | | | 73 | 5092 |
| 041A09963C | Female | LL | FS/WT | FS/L | | | | Mature | | | | | | | 76 | 5804 |
| 041A7192CB | Female | LL | FS/WT | FS/L | | | | Mature | | | | | | | | |
| 900119000414512 | Female | EL | FS/WT | FS/L | | | | Mature | | | | | | | 74 | 7528 |
| 900119000042107 | Female | EL | FS/WT | FS/L | | | | Mature | | | | | | | 82 | 6906 |
| 989001003400292 | Female | LL | FS/WT | FS/L | | | | Mature | | | | | | | 74 | 5382 |
| 989001003399347 | Female | LL | FS/WT | FS/L | | | | Mature | | | | | | | 82 | 9370 |
| 985170001826664 | Female | LL | FS/WT | FS/L | | | | Mature | | | | | | | 81 | 7592 |
| 041A7258A2 | Female | LL | FS/WT | FS/L | | | | Mature | | | | | | | 77 | 5676 |
| 041A5A81C0 | Female | LL | FS/WT | FS/L | | | | Mature | | | | | | | 82 | 7614 |
| 041A5B6E43 | Female | EL | FS/WT | FS/L | | | | Mature | | | | | | | 84 | 6490 |
| 041A09A1A7 | Female | EL | FS/WT | FS/L | | | | Mature | | | | | | | 78 | 6920 |
| 0006752139 | Female | LL | FS/WT | FS/L | | | | Mature | | | | | | | 82 | 7048 |
| 0419E5C947 | Female | EL | FS/WT | FS/L | | | | Mature | | | | | | | | |
| 041A7192D7 | Female | EL | FS/WT | FS/L | | | | Mature | | | | | | | 81 | 9010 |
| 992002000253861 | Female | EL | FS/WT | FS/L | | | | Mature | | | | | | | 89 | 10450 |
| 900119000342670 | Female | LL | FS/WT | FS/L | | | | Mature | | | | | | | 78 | 7876 |
| 900119000455645 | Female | LL | FS/WT | FS/L | | | | Mature | | | | | | | 78 | 8326 |
| 041A5B6C09 | Female | LL | FS/WT | FS/L | | | | Mature | | | | | | | 85 | 8636 |
| 900119000089164 | Female | EL | FS/WT | FS/L | | | | Mature | | | | | | | 86 | 10392 |
| 989001003400921 | Female | EL | FS/WT | FS/L | | | | Mature | | | | | | | 81 | 6992 |
| 900119000342082 | Female | EL | FS/WT | FS/L | | | | Mature | | | | | | | 82 | 6018 |
| 900119000088942 | Female | LL | FS/WT | FS/L | | | | Mature | | | | | | | 84 | 8922 |
| 0419E626F0 | Female | EL | FS/WT | FS/L | | | | Mature | | | | | | | 75 | 4924 |
| 041A711901 | Female | LL | FS/WT | FS/L | | | | Mature | | | | | | | 85 | 9312 |
| 900119000470414 | Female | EL | FS/WT | FS/L | | | | Mature | | | | | | | 79 | 8160 |

| | | | | | | | | | | | | |
|---|---|---|---|---|---|---|---|---|---|---|---|---|
| 04193FC6D4 | Female | EL | FS/WT | FS/L | | | Mature | | | | | |
| 985170001756945 | Female | EL | FS/WT | FS/L | | | Mature | | | | | |
| 041A723ACC | Female | EL | FS/WT | FS/L | | | Mature | | | | 55 | 2096 |
| 900119000342016 | Female | EL | FS/WT | FS/L | | | Mature | | | | 79.5 | 6788 |
| 900212000070781 | Female | LL | FS/WT | FS/L | | | Mature | | | | 79.5 | 8255 |
| 982000407903368 | Female | LL | FS/WT | FS/L | | | Mature | | | | 89 | 9957 |
| 900119000089727 | Female | LL | FS/WT | FS/L | | | Mature | | | | 82 | 7141 |
| 0419E605A2 | Female | EL | FS/WT | FS/L | | | Mature | | | | 76 | 7022 |
| 041A726001 | Female | EL | WT/WT | EL | | | Mature | | | | 89 | 11648 |
| 900119000342858 | Female | EL | WT/WT | EL | | | Mature | | | | 84 | 11806 |
| 900119000089876 | Female | EL | WT/WT | EL | | | Mature | | | | | |
| 900119000344422 | Female | EL | WT/WT | EL | | | Mature | | | | 86.5 | 10568 |
| 041A5AA9F4 | Female | EL | WT/WT | EL | | | Mature | | | | 79 | 6884 |
