## Supplementary File S2 for "Vgll3a promotes sexual maturation in male and female Atlantic salmon"

| PIT tag | Year-class | Sex | WT (%) | In-frame (%) | Frame-shift (%) | Total read count | Offspring year-class |
|---|---|---|---|---|---|---|---|
| 900119000341219 | 2016YC | Female | 12.21 | 11.29 | 76.51 | 33115 | 2020 year-class |
| 900119000341903 | 2016YC | Female | 9.04 | 22.89 | 68.07 | 45148 | 2020 year-class |
| 041A09A691 | 2017YC | Male | 5.82 | 14.78 | 79.4 | 34658 | 2020 year-class |
| 900119000343611 | 2017YC | Male | 14.72 | 9.25 | 76.03 | 41858 | 2020 year-class |
| 041A046D15 | 2017YC | Female | 0.02 | 18.63 | 81.36 | 37016 | 2021 year-class |
| 900119000343332 | 2017YC | Female | 7.36 | 11.35 | 81.28 | 35313 | 2021 year-class |
| 0419E63909 | 2017YC | Male | 6.7 | 12.25 | 81.06 | 33334 | 2021 year-class |
| 900119000341753 | 2017YC | Male | 1.33 | 12.53 | 86.15 | 33873 | 2021 year-class |
