## Supplementary File S3 for "Vgll3a promotes sexual maturation in male and female Atlantic salmon"

**Primers used in an PCR assay to genotype a large deletion of 442 bp (Del442) in vgll3a CRISPR target site.**

| Description | Sequence (5' - 3') | Amplicon size |
|---|---|---|
| Forward primer, up-stream of Del442 | CCTGGGTATCACAGAACTTG | |
| Reverse primer, downstream of Del442 | GCTCATCCACCACATCCTCG | WT: 753 bp, Del442: 311 bp |
| Reverse primer, overlapping Del442 junction | CTGTCGGACCTCAAATGAAC | Del442: 175 bp |

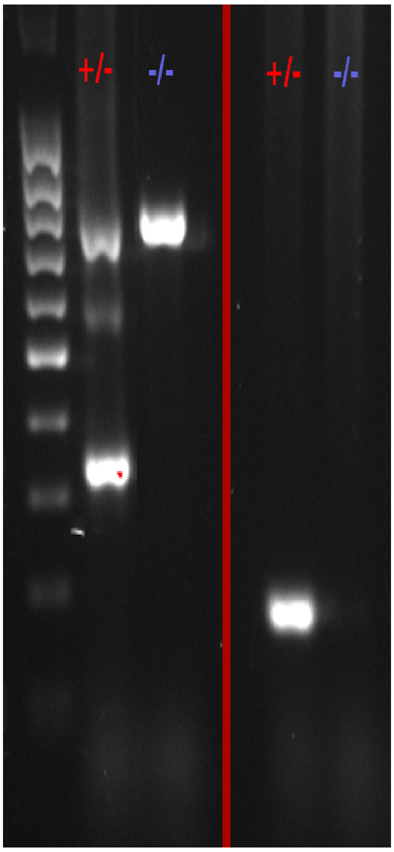


Description of figure: On the left, a Del442/WT (+/-) individual and a WT/WT (-/-) individual was assayed with upstream and downstream primers. On the right, the same individuals were assayed with upstream and junction primers.

**Primers and probes used in an allelic discrimination assay** to genotype a naturally occuring SNP giving amino acid replacement on aa 323 in *Vgll3a* linked to age at maturation. Sequences were obtained from Ayllon et al., 2019 (*BMC Genet*. 2019 May 6;20(1):44. doi: 10.1186/s12863-019-0745-9.).

| **Description** | **Sequence (5' - 3')** |
|---|---|
| Forward primer | AGCCCAGGGATACACAGTGA |
| Reverse primer | GTGGGCCAGGCTGAGG |
| Probe (early maturation) | CCACCTCTGT**C**TTCACA |
| Probe (late maturation) | CCACCTCTGT**G**TTCACA |

**Primers and probes used in qPCR assay to determine genetic sex** by detecting the presence of exons 2 and 4 of *sdy* gene. Protocol and original sequences were obtained from Ayllon et al., 2019 (*BMC Genet*. 2019 May 6;20(1):44. doi: 10.1186/s12863-019-0745-9.) but were here replaced with corrected sequences (Ayllon, personal communication). * indicates corrected sequences.

| Target | Primer/probe name | Sequence (5' - 3') |
|---|---|---|
| gapdh | Ss_gapdh_F | CCGCCACCCAGAAGACTGT |
| gapdh | Ss_gapdh_R | CTGGCGCCACGTCCAT |
| gapdh | Ss_gapdh_Pro | 6FAM-TCCTTCTGGAAAGCTGTGGA-MGBNFQ |
| sdy exon 2 | Ss_sdY_Exon2_F* | CCTACAAGCCCTTCTCCCTGAT |
| sdy exon 2 | Ss_sdY_Exon2_R* | GGGCTTTGGGAGAGAGATGAC |
| sdy exon 2 | Ss_sdY_Exon2_Pro* | VIC-ATGGATGGGATCCC-MGBNFQ |
| sdy exon 4 | Ss_sdY_Exon4_F | CCATGGGCTCAGCAGCTATT |
| sdy exon 4 | Ss_sdY_Exon4_R* | GGAGGACTCAAGCCAGATCCT |
| sdy exon 4 | Ss_sdY_Exon4_Pro | NED-AAGCAAGCTCACGACTT-MGBNFQ |
