## Supplementary File S4 for "Vgll3a promotes sexual maturation in male and female Atlantic salmon"

This section describes analysis of *vgll3a* isoform expression and linkage to E and L alleles in 10 immature testis samples from Atlantic salmon.

### Material and methods

Fertilization was performed 7th of November 2024, crossing two males and two females, producing four families that were mixed. The fish were kept on constant light in freshwater throughout the experiment. On the 29th of September 2025, fish were anesthetized with 100 mg/L Finquel vet. Body weight and gonad weight were measured. Fish with female gonads were discarded. Male gonads were stored on RNAlater (Thermo Fisher). GSI was calculated as (gonad weight / body weight) * 100%. None of the males were mature (GSI = 0.02-0.03). Fin clips were collected and used for DNA extraction on Biomek i5 Automated Workstation (Beckman Coulter). Genotyping of natural vgll3 alleles (E and L) was done with an allelic discrimination assay as described in the main text. RNA was extracted from 10 testis samples having EL genotypes and sequenced as 150 bp paired-end Illumina reads (Novogene). Sequence reads were mapped to the Atlantic salmon reference genome (Ssal_v3.1) using STAR (v. 2.7.11b, https://github.com/alexdobin/STAR). Mapped reads overlapping the start position of *vgll3* exon 2 were filtered (including their mates), using Samtools (v. 1.22.1, https://www.htslib.org/). This was done to ensure that the allele counts were derived from reads mapping to either the canonical or alternative isoforms. Ambiguously mapped reads, defined as reads where a single base was aligned on one side of the intron junction, or mapped to the intron sequence, were not included in the analysis. Read pairs mapped to either the canonical or alternative isoforms were counted. Read pairs containing natural alleles at a missense SNP at position 28,996,360 (E='T', L='C', causing a T54M amino acid substitution) were counted in reads from the canonical and alternative isoforms.

### Results

All of the 10 testes except one, had at least one read mapping to the alternative isoform (0-5 reads per individual), corresponding to 13% reads mapped to the alternative isoform, compared to 87% mapped to the canonical isoform, per individual on average (Table 1). Taking the alleles at the T54M missense SNP into account, on average, 1.2 and 1.5 reads mapped to the alternative isoform contained the E and L alleles, respectively (Figure 1). 8.3 and 7.1 reads mapped to the canonical isoform contained the E and L alleles, respectively (see Table 1 for read counts and Figure 1 for E and L allele counts associated with the alternative isoform). Mapping of all read pairs overlapping the exon 2 start site is shown for each sample in Figure 2.

### Discussion

We observed the same alternative isoform previously identified (Verta et al, 2020; https://doi.org/10.1371/journal.pgen.1009055). The study reported that the alternative isoform was associated with the allele for late maturation (L), however, we observe that the reads mapped to the alternative isoform which also overlap the T54M missense SNP contain the E and L alleles in similar proportions. This indicates that neither the E or L allele is associated with the alternative isoform.

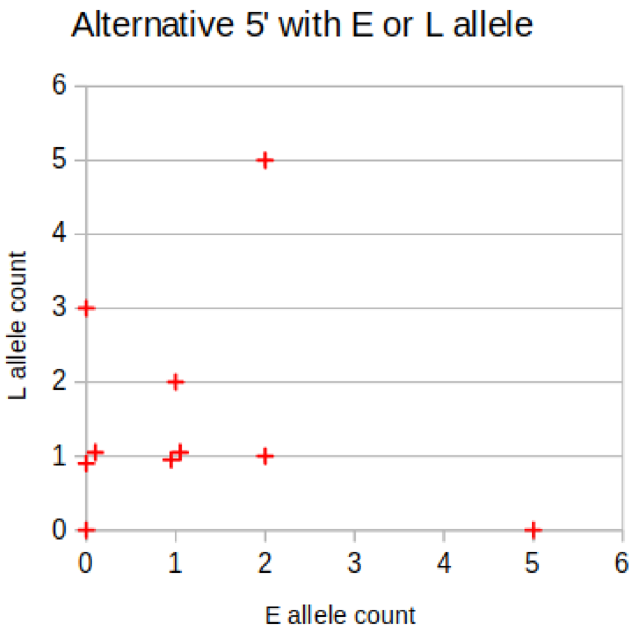


**Figure 1.** Count of read pairs mapping to the 5’ UTR of the alternative isoform, that also contained E or L alleles at the T54M missense SNP.

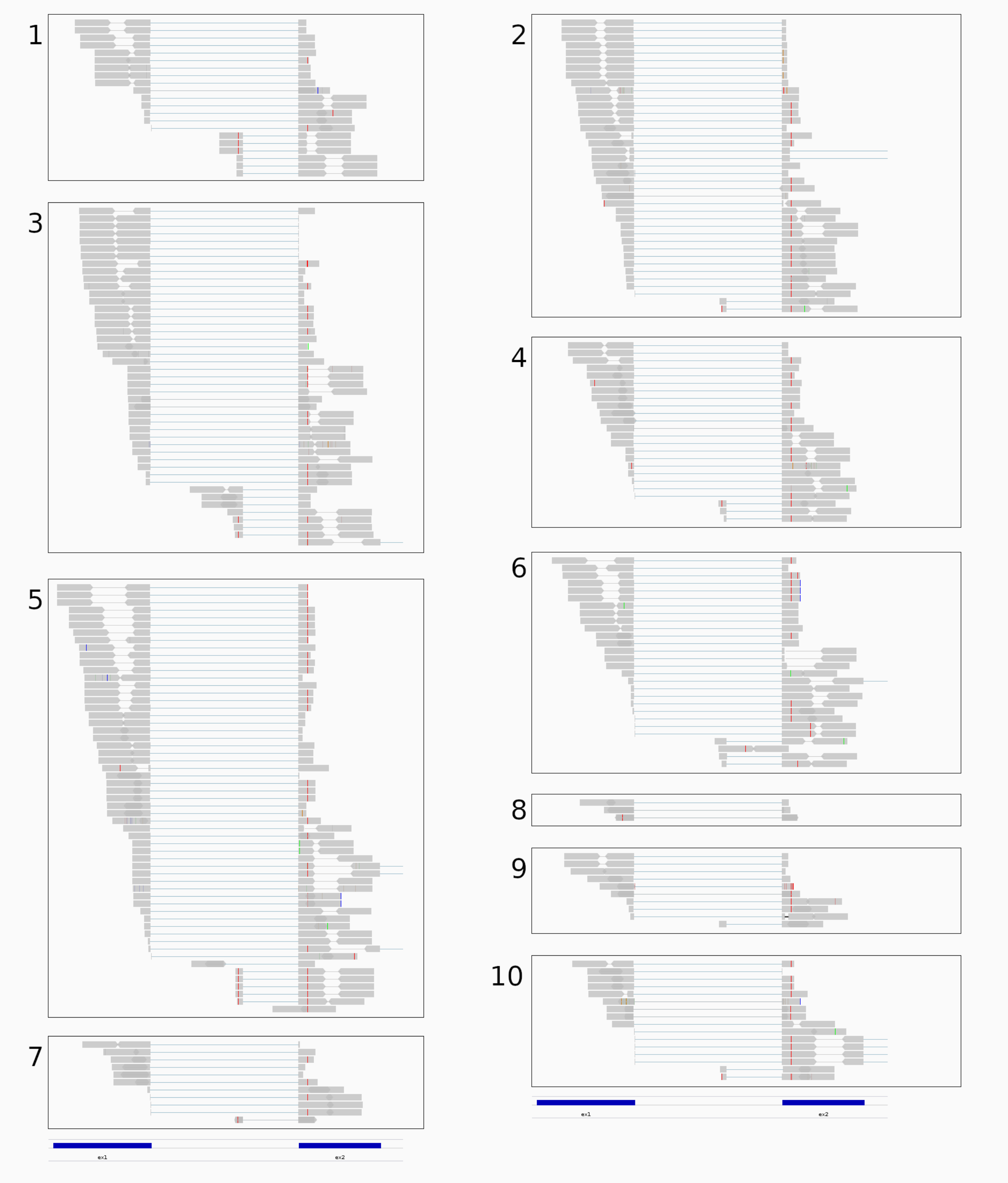


**Figure 2.** RNA-Seq reads mapped to *vgll3a*, overlapping with either canonical or alternative isoforms. Blue boxes at the bottom indicate locations of exon 1 and 2. The common red nucleotide in beginning of exon 2 represents the E-allele (L-allele in gray) of the T54M missense SNP. Numbers indicate sample number.

**Table 1.** Count of read pairs mapped to canonical and alternative isoforms, and reads mapped to either isoforms linked to E or L alleles at the T54M missense SNP. Reads with ambiguous mapping were not included in the analysis.

| **Sample** | **Canonical isoform** | **Alternative isoform** | **Ambigously mapped** | **Alternative + E** | **Alternative + L** | **Canonical + E** | **Canonical + L** |
|---|---|---|---|---|---|---|---|
| 1 | 14 | 6 | 1 | 0 | 3 | 1 | 11 |
| 2 | 36 | 2 | 1 | 1 | 1 | 16 | 6 |
| 3 | 31 | 7 | 7 | 2 | 5 | 13 | 12 |
| 4 | 19 | 3 | 2 | 2 | 1 | 8 | 9 |
| 5 | 48 | 5 | 4 | 5 | 0 | 24 | 16 |
| 6 | 21 | 3 | 4 | 1 | 2 | 8 | 9 |
| 7 | 8 | 1 | 2 | 0 | 1 | 3 | 2 |
| 8 | 3 | 0 | 0 | 0 | 0 | 0 | 2 |
| 9 | 9 | 1 | 0 | 0 | 1 | 4 | 2 |
| 10 | 8 | 2 | 6 | 1 | 1 | 6 | 2 |
